# Maize (*Zea mays L*.) nucleoskeletal proteins regulate nuclear envelope remodeling and function in stomatal complex development and pollen viability

**DOI:** 10.1101/2020.12.23.424208

**Authors:** JF McKenna, HK Gumber, ZM Turpin, AM Jalovec, AC Kartick, K Graumann, HW Bass

## Abstract

In eukaryotes, the nuclear envelope (NE) encloses chromatin and separates it from the rest of the cell. The Linker of Nucleoskeleton and Cytoskeleton (LINC) complex physically bridges across the NE, linking nuclear and cytoplasmic components. In plants, these LINC complexes are beginning to be ascribed roles in cellular and nuclear functions, including chromatin organization, regulation of nuclei shape and movement, and cell division. Homologs of core LINC components, KASH and SUN proteins, have previously been identified in maize. Here, we characterized the presumed LINC-associated maize nucleoskeletal proteins NCH1 and NCH2, homologs of members of the plant NMCP/CRWN family, and MKAKU41, homologous to AtKAKU4. All three proteins localized to the nuclear periphery when transiently and heterologously expressed as fluorescent protein fusions in *Nicotiana benthamiana*. Overexpression of MKAKU41 caused dramatic changes in the organization of the nuclear periphery, including nuclear invaginations that stained positive for non-nucleoplasmic markers of the inner and outer NE, and the ER. The severity of these invaginations was altered by changes in LINC connections and the actin cytoskeleton. In maize, MKAKU41 appeared to share genetic functions with other LINC components, including control of nuclei shape, stomatal complex development, and pollen viability. Overall, our data show that NCH1, NCH2, and MKAKU41 have characteristic properties of LINC-associated plant nucleoskeletal proteins, including interactions with NE components suggestive of functions at the nuclear periphery that impact the overall nuclear architecture.

## INTRODUCTION

In plant cells, as with all eukaryotes, the nucleus is a conspicuous and characteristic organelle, housing the DNA (reviewed by (Meier et al. 2017)). The nucleus itself is a dynamic structure, organized largely by its enclosing membrane system - the nuclear envelope (NE). The NE is a double-membraned structure composed of an outer nuclear membrane (ONM) and an inner nuclear membrane (INM) connected at nuclear pores (reviewed by (Hetzer 2010; Graumann and Evans 2017). The functional properties of the NE are mediated by protein complexes linked to myriad cellular and nuclear processes, including cell division and gene expression (Kim et al. 2015; Meier et al. 2017; De Magistris and Antonin 2018; Pradillo et al. 2019). A major conserved NE complex is the linker of nucleoskeleton and cytoskeleton (LINC) complex (Crisp et al. 2006), known for it involvement in processes such as the maintenance of nuclear architecture and mechanical structures, signalling, nuclear motility, and nuclear positioning (reviewed by (Rothballer and Kutay 2013), (Chang et al. 2015; Tamura et al. 2015; Pradillo et al. 2019; Starr 2019). Well beyond the historically recognized role of compartmentalization, NE investigations increasingly address how the NE contributes to both general and specialized functions in plant growth and development. The underlying molecular mechanisms coordinating and regulating these fundamental NE processes remain, however, largely unknown.

The hallmark of LINC complexes is their ability to form direct connections bridging the cytoplasm and the nucleoplasm. This is accomplished by a core complex of two groups of proteins, residing on the two separate membranes of the NE. The INM houses the Sad1/UNC84 homology (SUN) domain proteins and the ONM houses the Klarsicht/ANC-1/Syne homology (KASH) proteins (Hagan and Yanagida 1995; Malone et al. 1999; Starr 2002; Murphy et al. 2010). As a group, the KASH proteins are numerous and diverse, reflecting their various cytoplasmic binding partners, such as cytoskeletal and motor proteins (Starr and Fridolfsson 2010; Luxton and Starr 2014; Kim et al. 2015; Evans and Graumann 2018; Starr 2019). In contrast, the SUN-domain proteins are less diverse but still exhibit interactions with multiple components of the nucleoplasm, such as nucleoskeletal components and chromatin proteins (Haque et al. 2006; Janin et al. 2017). While cytoskeletal components of LINC complexes and the SUN proteins appear conserved in eukaryotes, plants have evolved unique KASH proteins and nucleoskeletal components. Plant nucleoskeletal components, while lacking sequence homology, share many of the animal lamin features, including interactions with the NE and their ability to impact chromatin structure and gene expression (Masuda et al. 1997; Dittmer et al. 2007; Wang et al. 2013; Ciska and Moreno Dí-az de la Espina 2014; Goto et al. 2014; Zhao et al. 2016; Guo et al. 2017; Choi et al. 2019; Ciska et al. 2019; Hu et al. 2019; Sakamoto 2020).

Plant nucleoskeletal proteins that interact directly or indirectly with the NE fall into two families of proteins. One family, the Nuclear Matrix Constituent Proteins/Crowded Nuclei (NMCP/CRWN) proteins, is found in plants with members encoded by the NMCP genes in carrot (Masuda), CRWN genes in Arabidopsis (Dittmer et al. 2007), and NCH genes in maize (Gumber et al. 2019a). The other family is also found in plants and encoded by the AtKAKU4 gene in Arabidopsis (Goto et al. 2014) and the MKAKU41 and MKAKU42 genes in maize (Gumber et al. 2019a). Here we refer to these two gene or protein families generally as CRWN and KAKU4.

In Arabidopsis, CRWN1 is located primarily at the nuclear periphery and interacts with SUN-domain proteins (Dittmer et al. 2007; Graumann 2014). CRWN proteins have been implicated in the regulation of nuclear shape, nuclear size, chromatin organization, regulation of gene expression, and nuclear body formation in plants (reviewed in (Sakamoto 2020). KAKU4 was shown to interact with CRWN1 by yeast two-hybrid analysis, supporting their cooperation in the form and function of a plant nucleoskeletal system (Goto et al. 2014). Fluorescent protein fusions driven by native promoter expression and electron microscopy showed that KAKU4 is localized to the inner nuclear membrane. Interestingly, when high-expression stable lines were selected or high-expression transient transformation was performed, KAKU4 appeared to induce nuclear invaginations, which increased significantly when co-expressed with CRWN1 (Goto et al. 2014).

Together, the CRWN and KAKU proteins are known to affect phenotypes in eudicots, yet their roles in crop species are less well understood. Two maize CRWN homologs are known; NCH1 which is most closely related to AtCRWN1-3, and NCH2 which is most closely AtCRWN4. Maize MKAKU41 homologs are encoded by *MKAKU41* and *MKAKU42* (Gumber et al. 2019a). To investigate the functional conservation of these presumed nucleoskeletal proteins in maize, we characterized *NCH1* and *NCH2* and *MKAKU4* using cytological or genetic approaches.

## RESULTS

### Maize NCH1, NCH2, and MKAKU41 localized to the nucleus, primarily at the nuclear periphery

In order to determine the cellular localization of the maize CRWN homologs NCH1, NCH2, and the KAKU4 homolog MKAKU41, we produced gene constructs with the protein coding region fused to either GFP or mCherry at the N-terminus. We then expressed these constructs transiently in *N. benthamiana* leaf tissue. All three constructs localized to the nuclear periphery with MKAKU41 also exhibiting internal structures as shown in Figure 1 for nuclei counterstained with DAPI. In order to confirm that NCH1 and NCH2 were localized at the nuclear periphery, we performed co-expression of NCH1 or NCH2 with AtCRWN1 (Fig. S1). The co-localization of NCH with CRWN confirmed that NCH1 and NCH2 did localize to the nuclear periphery when transiently expressed in *N. benthamiana*. Interestingly, MKAKU41 showed nuclear envelope labelling and inner nucleus labelling in most cells, including membrane invaginations (Fig. 1B, white arrowheads), as has been described for Arabidopsis KAKU4 when expressed under the control of the 35S promoter (Goto et al. 2014). These included internal structures and circular ring-like invaginations within the nucleus. The structures labelled with MKAKU41 colocalized with brighter DAPI staining regions, implying that MKAKU41 may associate with heterochromatin. The ring-like invaginations lacked internal DAPI staining and therefore appeared devoid of typical nucleoplasmic chromatin. Interestingly, the CRWN1 homolog NCH1 alone also caused these aberrant-looking intranuclear structures (Fig. 1B, white arrow), although to a lesser extent than MKAKU41. However, NCH2, a CRWN4 homolog (Fig. 1A), did not induce ring-like invaginations. The NCH1 and NCH2 also displayed different mobility proportions at the nuclear periphery (Fig. 1C, S1B and S1C) as determined by Fluorescence Recovery After Photobleaching (FRAP). NCH2 was significantly less mobile (16% mobile fraction) than NCH1 (49% mobility) indicating that they might interact with different protein complexes or structures. The lower mobility of NCH2 compared to NCH1 is consistent with its reduced capacity to remodel the nuclear envelope membrane and cause invaginations (Fig. 1A,C). The large immobile fraction of NCH2, but not NCH1, was comparable to that of AtCRWN1 (Graumann 2014).

**Figure 1:**
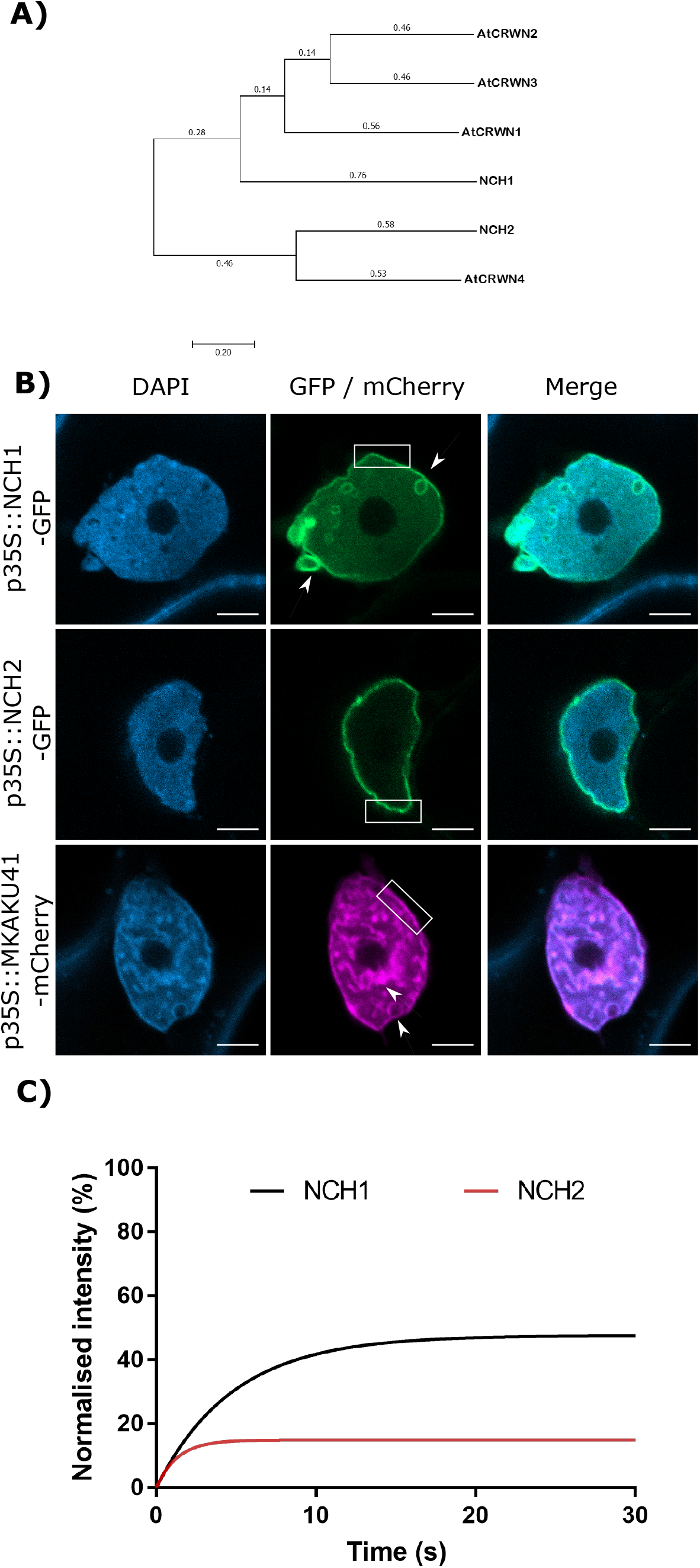
The maize proteins NCH1, NCH2 and MKAKU41 localize to the nuclear envelope. (A) Maximum likelihood phylogenetic tree showing maize NCH1 and NCH2 along with homologous proteins CRWN1-4 from Arabidopsis. (B) Confocal imaging of *N. benthamiana* transiently expressing NCH1, NCH2 and MKAKU41 as fluorescent protein fusions (green / magenta) and nuclei labelled with DAPI (blue). Both MKAKU41 and NCH1 show nuclear invaginations (white arrowheads) and labelling at the nuclear envelope (white boxes). NCH2 shows predominantly nuclear peripheral localization. Scale bar denotes 5μm. (C) FRAP analysis demonstrates that NCH1 has a larger mobile fraction than NCH2 at the nuclear periphery. N ≥ 30 nuclei imaged across three experimental replicates.

It has been demonstrated that the degree of nuclear invagination and deformation is dependent on the expression level of KAKU4 in *A. thaliana* (Goto et al. 2014). We therefore asked if this causal relationship was conserved in the maize genes, MKAKU41, NCH1, and NCH2, using the dose-response transient expression assays summarized in Figure 2. We infiltrated *N. benthamiana* with three different concentrations of Agrobacterium, OD 0.01, 0.05, or 0.1, and then classified live-imaged nuclei (N≥30 per condition) into one of three previously described nuclear periphery cytological image patterns (Goto et al. 2014): type I for normal nuclear periphery localization, type II for minor invagintions or inclusions in the nucleus, and type III for major invaginations and deformation of the nucleus. This classification assisted with a comparative analysis of the severity of changes in nuclear morphology across the various experiments used throughout this study. Increasing the infiltration concentration of NCH1 resulted in progressively increased nuclear deformation illustrated by type II pattern increases and the appearance of type III at the highest transfection dose (Fig. 2A). NCH2 also showed a transfection dose-dependent increase in nuclear aberration coupled to type II increases, but less so and without any type III nuclei even at the highest dose (Fig. 2B). For MKAKU41, increasing the transfecting plasmid concentration led from most nuclei just showing peripheral localization initially to slightly under half showing type III patterns with severe invaginations seen at the highest concentration (Fig. 2C). We interpret these results as establishing that because all three of the proteins tested showed they could cause dose-dependent increases in severity of nuclear deformation patterns, they can all be considered to be part of the same process, one needed to properly organize a nuclear periphery compartment within interphase nuclei.

**Figure 2:**
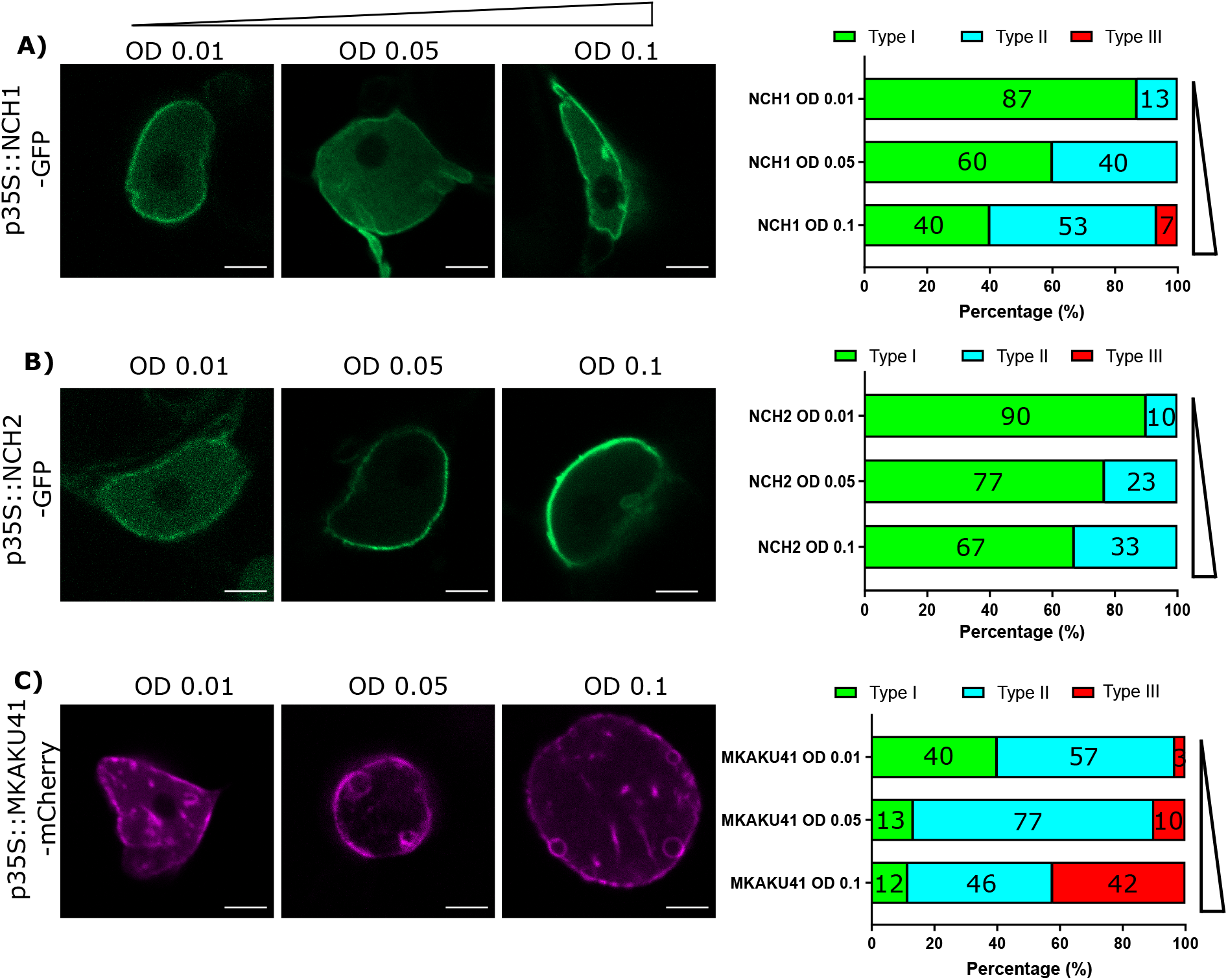
The concentration of NCH1, NCH2 and MKAKU41 affects the level of nuclei invagination and disruption. Confocal live cell imaging of NCH1, NCH2 and MKAKU41 shows that increasing the concentration of NCH1 and MKAKU41 increases nuclei disruption. Nuclei were classified as per (Goto et al. 2014) into Type I, Normal nuclear envelope localization; Type II, Minor invaginations and predominantly nuclear envelope localization; or Type III, major disruption of nuclei structure. (A) NCH1, (B) NCH2 and (C) MKAKU41. Scale bar denotes 5μm. N≥ 30 nuclei imaged across three experimental replicates.

### Co-expression of NCH1 or NCH2 with MKAKU41 showed a synergistic effect on invagination and nuclear deformation phenotypes

The Arabidopsis CRWN1 and KAKU4 have previously been shown to result in increased nuclear deformation and invaginations when co-expressed (Goto et al. 2014). In order to determine whether the maize homologs similarly affect the nuclear organization pathway, we co-expressed MKAKU41 with either NCH1 or NCH2 as presented in Figure 3. Upon co-expression, the nuclear invaginations and deformations were substantially enhanced compared to single-expression (Fig. 3A). Quantifying the most severe phenotype (type III), we observed that NCH1 alone exhibited 7% and MKAKU41 alone exhibited 42%. However, the combination of NCH1 and MKAKU41 exhibited 88% type III, well beyond the combined value of 49%, and thus synergistic for this measurement. Similarly, the NCH2 and MKAKU41 co-expression reached a level of 66% type III nuclei, considerably more than 42% sum of the single-expressed levels. Therefore, NCH proteins appeared to act cooperatively with MKAKU41 to affect nuclear periphery organization as judged by changes in their localization patterns. This cumulative effect on nuclear structure suggests that NCH1/NCH2 and MKAKU41 may act in the same protein complex. We tested this idea by checking for evidence of interaction between NCH and MKAKU41 *in planta* using acceptor photobleaching fluorescence resonance energy transfer (apFRET). A significant rise in FRET efficiencies would indicate interactions and such was measured when NCH1 and NCH2 were co-expressed with MKAKU41, compared to a non-interacting control, calnexin (Fig. 3C). Internal control FRET efficiency % values were very low, as expected for non bleached controls (Table S1). This demonstrated that NCH1 and NCH2 can interact with MKAKU41, and is consistent with their synergistic effect on levels of nuclear deformations. This observation is similar to that observed for Arabidopsis homologs using yeast two-hybrid system and plant expression (Goto et al. 2014). Importantly, our findings provide evidence from live imaging for *in planta* interactions between these proteins at the nuclear periphery.

**Figure 3:**
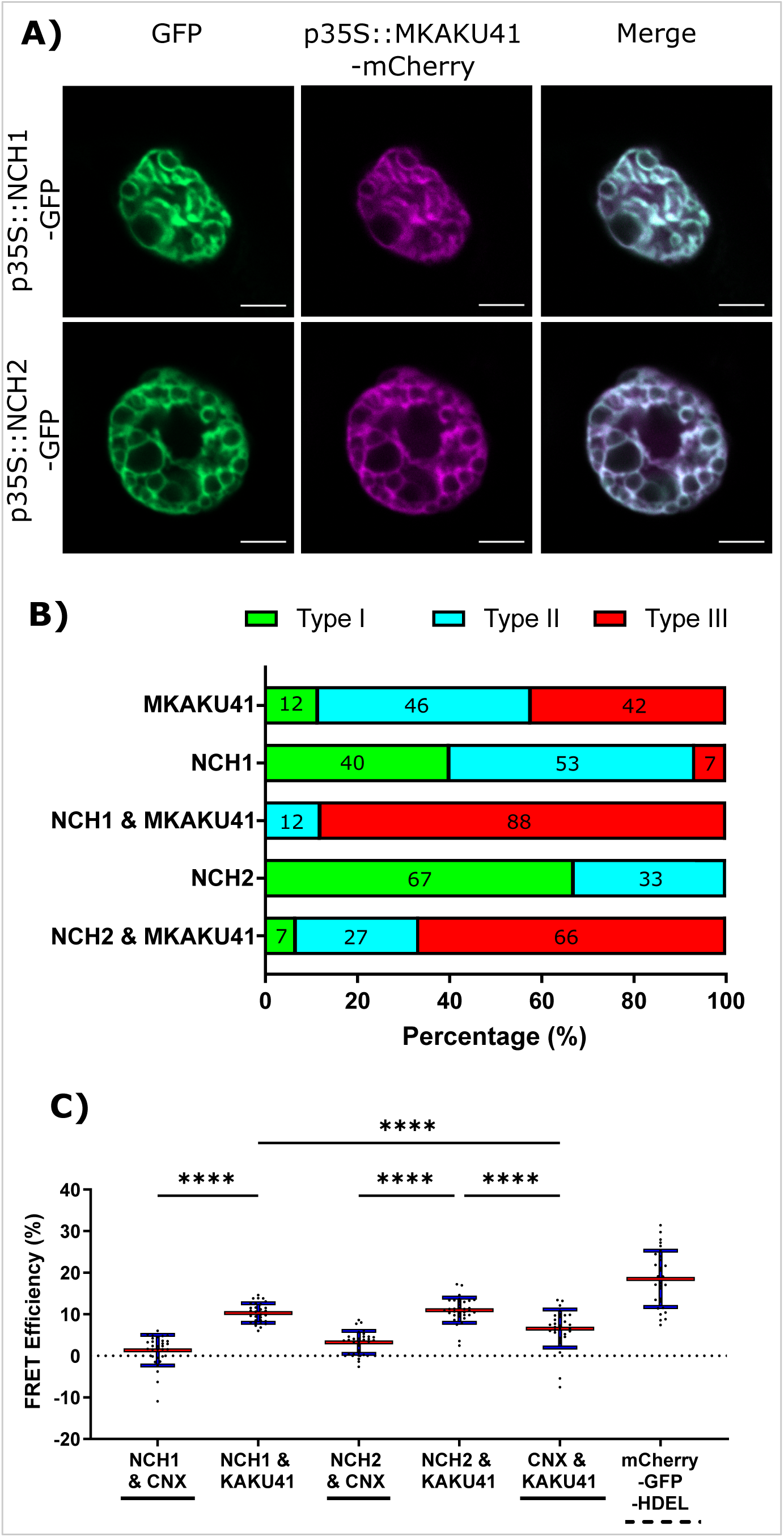
Coexpression of NCH1 or NCH2 with MKAKU41 enhances nuclear invaginations and NCH1 and NCH2 interact with MKAKU41. (A) Confocal images of N. benthamiana coexpressing NCH1 or NCH2 with MKAKU41. (B) Quantification of nuclear morphologies for nuclei expressing single MKAKU41, NCH1, NCH2; or co-expression of NCH1 or NCH2 with MKAKU41; classified into types I, II, or III as described in Figure 2. Scale bar denotes 5μm. (C) The apFRET efficiency (%) demonstrates in planta interactions of NCH1 and NCH2 with MKAKU41 in comparison with control (CNX). Solid lines underneath plots denote negative controls, hashed positive control. Key statistical comparisons shown and more fully tabulated in Table S1. Red line denotes mean, blue standard deviation. One-way ANOVA statistical test performed. **** = P ≤ 0.0001. N≥ 30 nuclei imaged across three experimental replicates for all experiments described.

### MKAKU41 overexpression remodeled other inner and outer nuclear membrane proteins

Both AtKAKU4 and MKAKU41 cause deformation of the NE and disruption of nuclei structure when co-expressed with AtCRWN1 (Goto et al. 2014), NCH1, or NCH2 (Fig. 3). To further investigate whether this involves the entire NE, we asked whether overexpression of MKAKU41 can result in the invagination of other proteins not previously tested but known to localize to either the INM, the ONM, or the ER. For this we used AtSUN2-YFP to mark the INM, ZmMLKP1-GFP to mark the ONM, and calnexin-GFP to mark the ER membrane. Figure 4 shows that each of these markers showed normal nuclear periphery localization when expressed individually. However, upon co-expression with MKAKU41, all of these proteins appeared in aberrant intranuclear structures. Therefore, the expression of MKAKU41 appears to have caused the internalisation of the entire NE, including the calnexin-GFP ER membrane marker, demonstrating that MKAKU41-induced nuclei deformations and invaginations are not limited to nucleoplasmic and INM LINC proteins.

**Figure 4:**
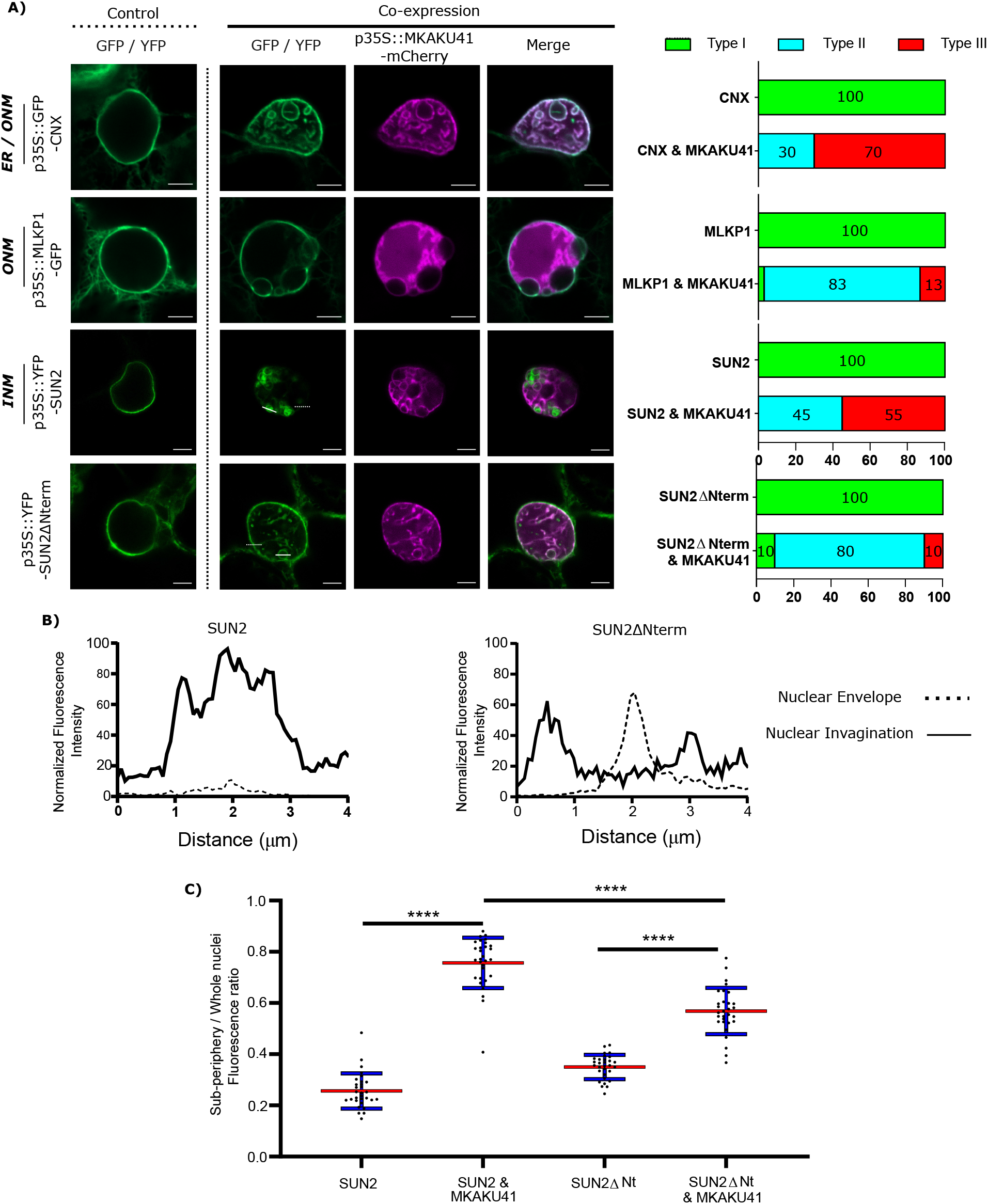
MKAKU41-induced nuclear invaginations incorporate other nuclear envelope- and ER-localized proteins. (A) Confocal imaging showing that the outer nuclear Maize membrane markers MLKP1, Arabidopsis nuclear envelope proteins SUN2 and SUN2ΔNterm, and the ER membrane marker CNX are incorporated into invaginations when co-expressed with MKAKU41. Next to confocal images are percentages of nuclei which show deformations as classified previously. (B) SUN2 and SUN2ΔNterm show different incorporation into MKAKU41 induced nuclear invaginations. Graphs show 4μm normalised line profiles over SUN2 or SUN2ΔNterm Invaginations (solid white lines) and nuclear membrane (dotted white lines). Locations of line profiles can be seen in confocal micrographs. (C) Ratio of Sub-periphery/whole nuclei fluorescence of SUN2 and SUN2ΔNterm when expressed on their own or with MKAKU41. **** = P ≤ 0.0001. N ≥ 30 for each condition.

Previously, it has been shown that AtSUN1 and AtSUN2 interact with AtCRWN1, mediated by the SUN N-terminus (Graumann 2014) and that the deletion construct SUN2ΔNterm could disrupt SUN-CRWN interactions (Graumann et al. 2010; Graumann 2014). To investigate more specifically the role of the LINC complex in producing these aberrant nuclear structures, we overexpressed SUN2ΔNterm as a way to abrogate CRWN-SUN binding and thereby possibly reduce the interaction of MKAKU41 with the LINC complex. Notably, SUN2-MKAKU41 co-expression resulted in mostly type III nuclei, whereas the SUN2ΔNterm-MKAKU41 co-expression led to less deformation, with nuclei showing mostly type II phenotype (Fig. 4A). Interestingly, nuclear envelope fluorescent labelling of full-length SUN2 appeared lower than that in SUN2ΔNterm when co-expressed with MKAKU41 (Fig. 4A & 4B). Signal intensity line profiles drawn over the nuclear invagination (Fig. 4A, dashed line) or nuclear envelope (Fig. 4A, solid line) regions showed that the SUN2 full length signal was much stronger within the invaginations but lower at the nuclear envelope when compared to those from SUN2ΔNterm fluorescence (Fig. 4B). We quantified the effect of co-expressed MKAKU41 on the SUN2’s tendency for peripheral staining by determining the percentage of the signal in the interior versus the entire nucleus (sub-periphery / whole nucleus/, Fig. S2). For instance, if the signal was entirely peripheral, the percentage would be at or near zero. Figure 4C shows an increase in internal fluorescence for SUN2 in the SUN2 - MKAKU41 co-expression compared to SUN2 alone (p<0.0001). Therefore, full length SUN2 is relocated away from the nuclear periphery when coexpressed with MKAKU41. The relative internal nuclear fluorescence also increased upon SUN2ΔNterm coexpression with MKAKU41 (p<0.001), but to a lesser extent than for full length SUN2 (p<0.001). These observations implicate the LINC complex and more specifically the nucleoplasmic N-terminal domain of the NE protein, SUN2, in the formation of aberrant MKAKU41-dependent intranuclear structures.

### Impairing Nuclear Actin anchoring enhances nuclear invaginations

In addition to the SUN2 NE protein which spans the INM and has direct contact with the nucleoplasm, we also examined an ONM maker, MLKS2, and its ARM domain deletion derivative previously shown to be impaired for actin binding (Gumber et al. 2019b). Upon co-expression with MKAKU41, the ONM marker MLKS2 also appeared in intranuclear invagination-like structures as shown in Figure 5 (panel A). Upon co-expression of MKAKU4 and MLKS2ΔARM, nuclear deformations and invaginations were observed (Fig. 5A) and found to much more severe and abundant (Fig. 5B) compared to those from co-expression of MKAKU4 and the full length MLKS2. Therefore, the ONM KASH protein MLKS2 is brought moved to intranuclear structures by MKAKU41 co-expression, and loss of the actin-interacting ARM domain exacerbates the situation and implicates cytoplasmic F-actin in NE and nuclear periphery remodeling.

**Figure 5:**
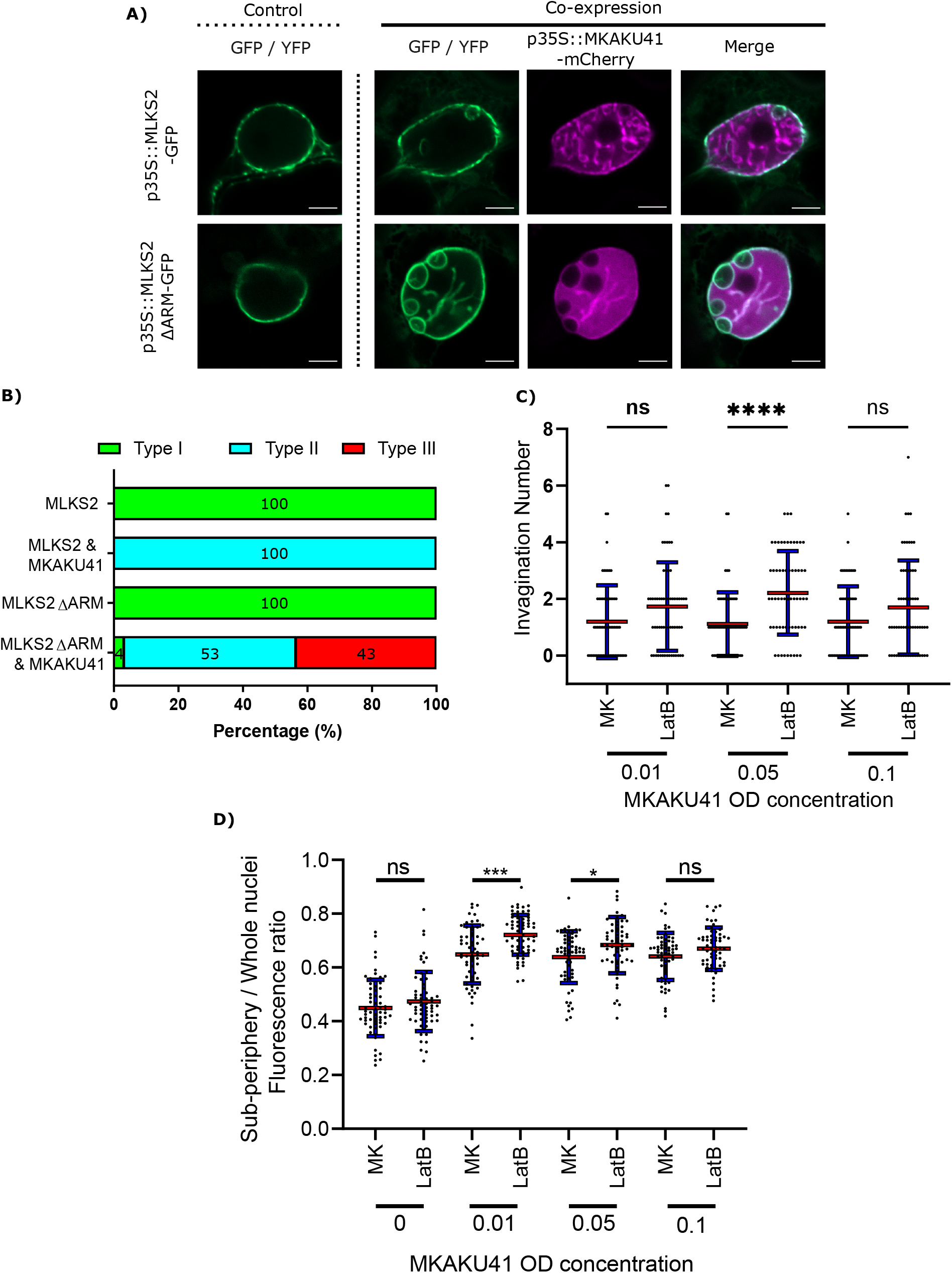
Nuclear actin anchoring is important for regulating nuclei deformations upon MKAKU41 overexpression. (A) Confocal live cell imaging of MLKS2 and MLKS2ΔARM single and co-expresssion with MKAKU41. (B) All nuclei imaged were categorised on the level of disruption as described previously. (C) Number of nuclei invaginations at different MKAKU41 expression levels with actin depolymerised (LatB). (D) Sub-periphery / whole nuclei fluorescence ratio from nuclei expressing different levels of MKAKU41 (MK, from 0 to 0.1) with and without actin depolymerisation (LatB). Nuclei were imaged with the NE marker LBR-GFP. Scale bar denotes 5μm. N≥ 30 nuclei imaged across three experimental replicates for panel A & B). For Panel C & D ≥ 60 across three biological replicates.

In order to further probe the interaction between peri-nuclear actin and MKAKU41-induced nuclear deformations, we used Latrunculin-B (LatB) to depolymerise the actin cytoskeleton as described previously (McKenna et al. 2019). Figure 5C shows that upon expression of MKAKU41 at a moderate level (OD0.05), LatB depolymerisation of the actin cytoskeleton resulted in a statistically significant increase in the number of invaginations (P≤0.001). While this trend existed at lower (OD0.01) and higher transfection concentrations (OD0.1), it was not statistically significant. To further explore the connection between the actin cytoskeleton and MKAKU41 induced nuclear deformations, we examined the interior versus peripheral signal ratios (Fig. S2) and found that actin depolymerization quantitatively shifted the INM NE marker, LBR, towards increased interior signal (Fig. 5D). This same effect was seen and found to be statistically significant for two concentrations of MKAKU41 infiltration, OD0.01 (P≤0.01) and 0.05 (P≤0.05). This demonstrates that at the moderate transfection concentration of 0.05OD, actin depolymerization with LatB increases the internalization of peripheral markers. These findings corroborate those from the MLKS2ΔARM experiments in that they implicate F-actin as a possible factor that can provide an opposing force or counterbalance to nucleoskeletal proteins which can invaginate the NE and create inclusion bodies in a concentration-dependent manner.

### Transposon disruption of the MKAKU41 gene co-segregates with phenotypic effects on nuclear shape and development

Having seen that overexpression of maize MKAKU41 in a eudicot species resulted in nuclear architecture disruption and severe nuclear envelope misplacement, we wanted to examine the role of this gene in its native genetic background, maize. From the UniformMu transposon mutagenesis project, we found and characterized a Mutator-tagged allele of *MKAKU41*, allowing for a genetic examination of the biological consequences of gene disruption in maize.

The transposon-tagged allele, here designated *mkaku41*, and its wild-type counterpart, *MKAKU41*, are shown in Figure 6. The wild-type *MKAKU41* gene (Zm00004b040444) from the color-converted W22 inbred is annotated as being associated with three transcript models. The gene structure for transcript model T01 (Fig. 6A) spans 7.9 kb, with 11 exons producing an mRNA with a single large ORF predicted to encode a protein of 579 AA. The transposon insertion site (mu1005806) is located in the 5’ UTR (Fig. 6B). The transposon insertion allele was characterized by genomic PCR analysis (Fig. 6C) using various combinations of primers that were flanking the insertion site, and were gene-specific and *Mutator*-specific (Fig. 6A, primers F, T, R). These PCR products were visualized (Fig. 6C) and sequenced (Fig. 6D) from the W22 wild type progenitor (W22+) as well as F2 individuals from families segregating for *mkaku41* (+/+, +/−, or −/−), where the “−” symbol denotes the transposon-insertion allele. These PCR primers and PCR gel products were used for plant genotyping in subsequent analyses.

**Figure 6.**
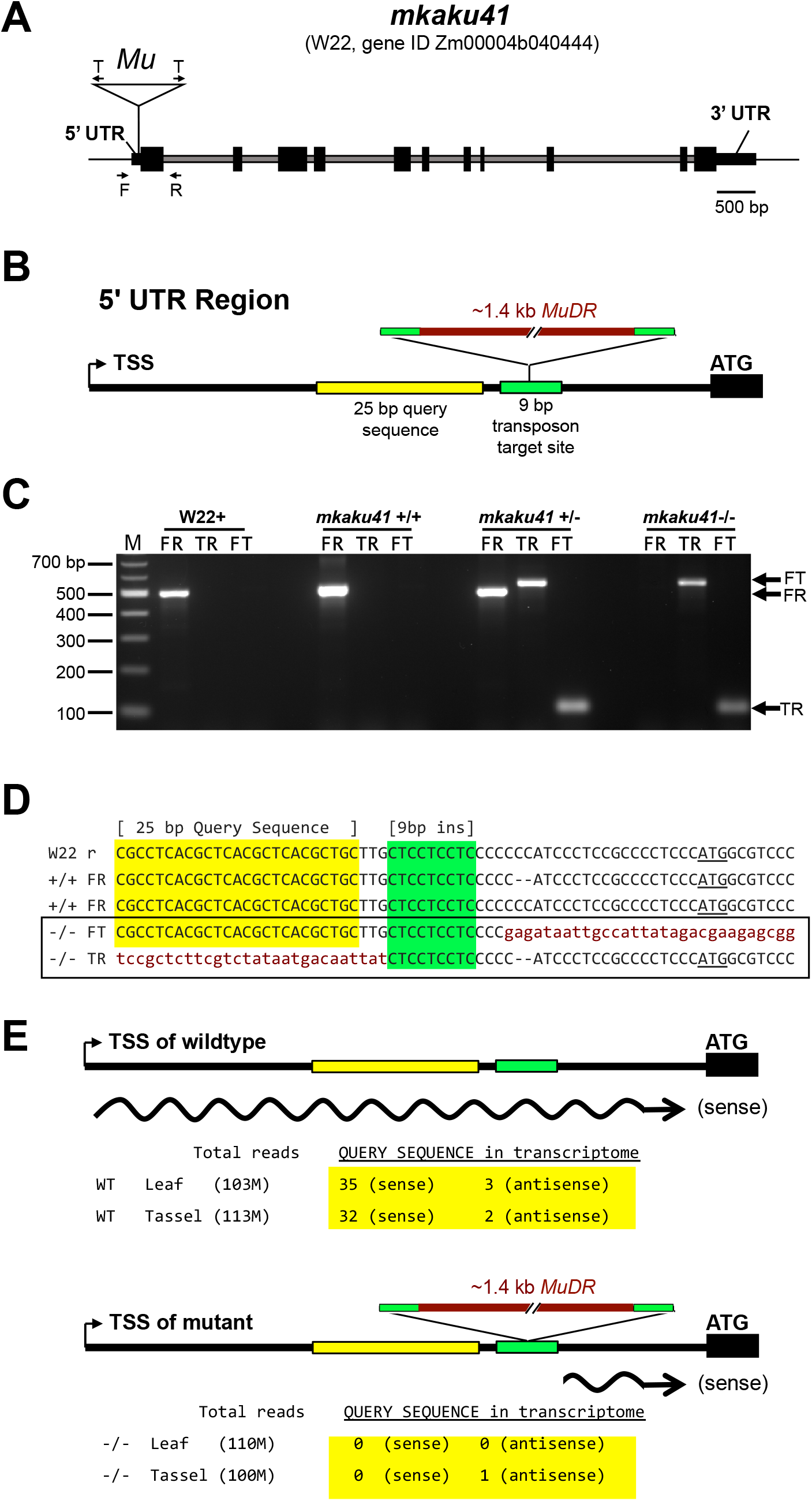
Gene structure of wild-type and transposon-tagged alleles of *MKAKU41*. (A) Gene model of MKAKU41 (transcript model “T01”) showing the positions of exons (black boxes), introns (grey), the 5’ and 3’ UTRs, the transposon-insertion (Mu), transposon-specific Tir6 PCR primers (T arrows), and the Mu-flanking gene-specific PCR primers (F, R arrows). (B) Diagram of the MuDR insertion site within the 95 bp 5’ UTR, showing the locations of the 9 bp target site repeat (green, bp 64-72 in the mkaku41-T01 transcript model) and a 25 bp query sequence (yellow) used for transcript analysis. (C) PCR Genotyping for presence or absence of the Mu-tagged allele. The PCR products using various pairings of gene-specific (F, R) or transposon-specific primer pairs (T). Ethidium bromide-stained agarose gel 100 bp size marker (lane M) and single-plant PCR products (other lanes) from W22+ or select mkaku41 F2 siblings to illustrate wildtype (+/+), heterozygous (+/−), or mutant (−/−) PCR genotype patterns from the PCR (arrows at right of gel). (D) Sequence of cloned and sequenced amplicons are aligned and show the progenitor W22 (W22 reference) and F2 segregants. The +/+ individuals from the transposon mutagenesis stocks were found to be heterozygous for a 2 bp indel (−−). The sequences corresponding to a 25 bp query sequence (yellow highlight), the 9 bp insertion site (green highlight), the transposon (lowercase, garnet text), and the start codon (underlined ATG) are indicated. The Mu insertion occured in the allele with the 2bp deletion and produced a flanking 9bp target site duplication. (E) Strand-specific transcriptome analysis is summarized for perfect match occurrences of the query sequence in libraries made from wildtype (top) or mutant (bottom) plants. All of the sense transcripts from the mutant allele had an extremely short (~21 bp or less) 5’UTR.

By inspection of the junction sequences between the wild-type reference W22 genome and the transposon (Fig. 6D, lower case letters), we identified the 9-bp target site duplication as CTCCTCTC (color coded green in Fig. 6). These results confirmed that the transposon was inserted in the 5’ UTR at a position 23 bp upstream of the start codon, disrupting the majority of the wild-type 95 bp 5’ UTR. The location of this insertion, while not in the protein coding region of the gene, is within the first exon and its location is expected, therefore, to disrupt the expression or transcript structure of the gene. Surprisingly, from transcriptome analysis we found that the gene expression levels for MKAKU41, measured as total normalized reads across the gene model, were similar in libraries made from wildtype and mutant leaf and tassel. We mined the transcript data to investigate the effect of the transposon insertion on the 5’ UTR region by searching for the presence of a unique 25 bp 5’ UTR sequence located just upstream of the mapped insertion site (Fig. 6D,E, “Query sequence” highlighted in yellow). We found a total of 70 matches to our 25 bp query sequence in our transcriptome, which was sequenced at a depth of over 100M reads per tissue-genotype combination (Fig. 6E). All 70 occurrences of the query sequence were from the wildtype libraries except for one, which was on the reverse strand relative to the gene (Fig. 6E). These results indicate that the 5’ UTR was indeed disrupted in the mutant plants. In addition to this detailed analysis of MKAKU41, some differentially expressed genes (Table S2) were observed in mutant versus wild-type leaf and meiotic-enriched whole tassel, but gene ontology analysis did not reveal any clear and reproducible enrichments that differed from those of randomized controls.

We next explored the phenotypic consequences of the *mkaku41* transposon insertion on root hair nuclei, stomatal complex, and pollen viability, as summarized in Figure 7. The root hair nuclei in W22+ (normal) and *mkaku41* mutant seedlings 5 days after imbibition were imaged and their shapes were analyzed (Fig. 7A-F). The mutant nuclei were visually and quantitatively more rounded than their wildtype counterparts. The mutant nuclei had an average maximum length of 22 μm whereas their wild-type counterparts averaged 34 μm (Fig. 7G). The mutant nuclei also exhibited a higher circularity index than the wildtype nuclei (Fig. 7H). Both measures (n=50) were statistically significant as determined using T-test, two-tailed with p<0.0001.

**Figure 7.**
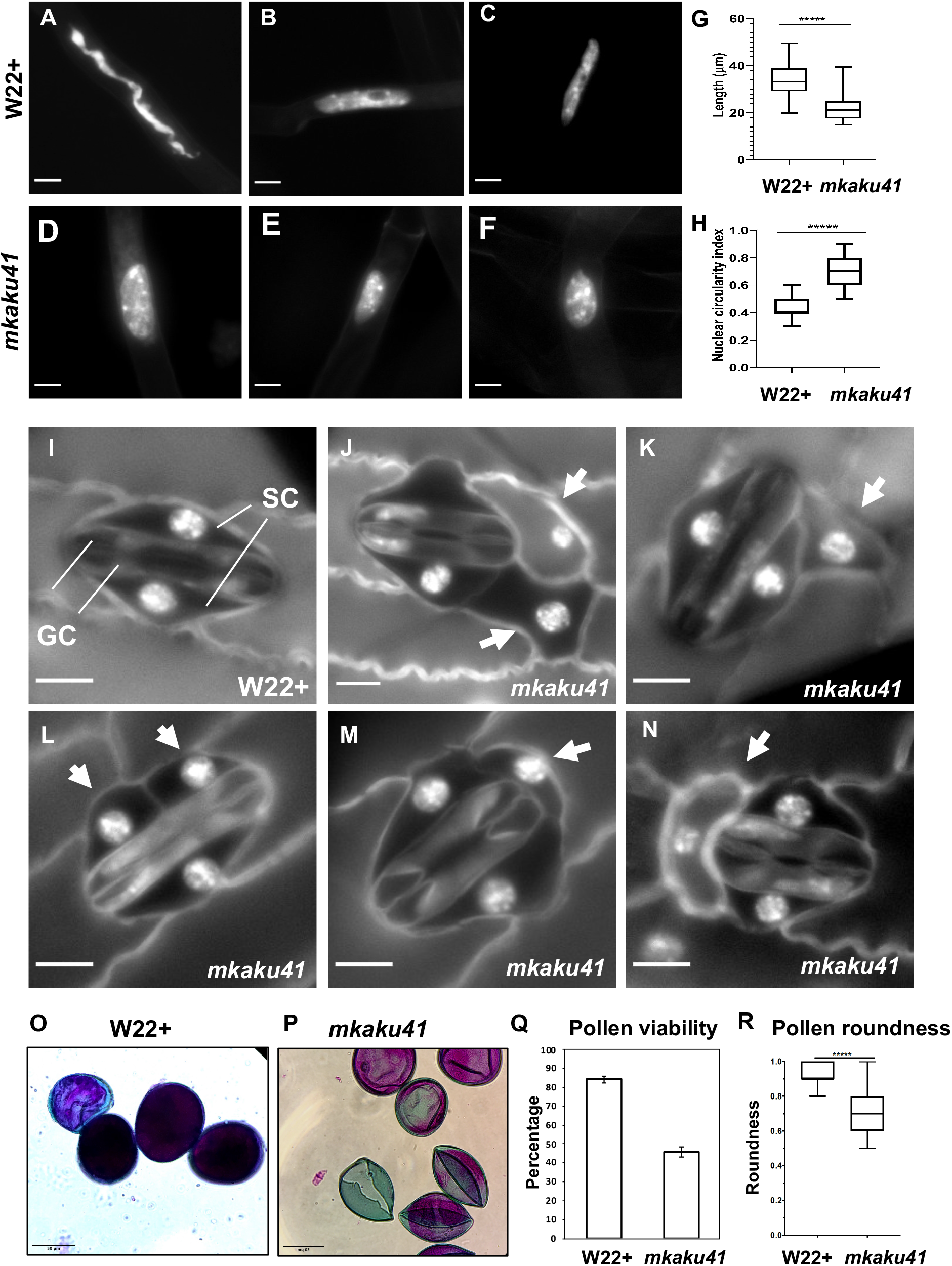
Somatic phenotypes of *mkaku41*. DAPI-stained root hair nuclei in W22+ (A-C) and mkaku41 (D-F) seedlings. (G) Longest diameter of W22+ and mkaku41 root hair nuclei (n=50). (H) Nuclear circularity index measurements using 4π(area/perimeter^2^), where 1= perfect circle, of W22+ and mkaku41 root hair nuclei calculated using Fiji. (I) Mature stomatal complex in W22+ DAPI-stained leaf where two central dumbbell-shaped guard cells (GC) are surrounded by two subsidiary cells (SC). (J-N) Representative images of stomatal complexes in *mkaku41* DAPI-stained leaves. Arrows point to extra or irregularly-placed subsidiary cells. Scale bars are 10 μm. *****Student’s t-test two-tailed, p < 0.0001. (O-Q) Differential staining of anthers for testing pollen viability from W22+ (O) and mkaku41 −/− (P,Q) tassels stained with modified Alexander’s stain where viable pollen grains appear magenta and aborted pollen grains appear green. (R) Quantification of pollen viability, n>1000 per genotype. (S) Degree of pollen roundness 4*area/(π*major_axis^2) calculated for W22+ and mkaku41−/− anthers using Fiji. Scale bars are 50 μm. *****Student’s t-test two-tailed, p < 0.0001.

Next, we analyzed two above-ground phenotypes, the appearances of stomatal complexes and pollen. In W22+ plants, a normal stomatal complex is composed of two guard cells flanked by two subsidiary cells as shown for W22+ (Fig. 7I). In contrast, mutant plants showed irregular stomatal complexes composed of two normal-looking guard cells flanked by one or two extra and irregularly positioned subsidiary cells (arrows, Fig.7 J-N). For the pollen phenotypes, we assayed viability and shape (Fig. 7O-R). Using the modified Alexander’s differential staining method, we found that the percent of viable pollen was dramatically reduced in the mutant, from 84% to 46%. The shape of the pollen was also affected in the mutant, where the average degree of roundness decreased from 0.93 to 0.7. Both measures (n>1,000 for staining, n=100 for roundness) were statistically significant as determined using a two-tailed T-test with p<0.0001.

Taken together, these findings show that the *mkaku41* mutation was associated with multiple phenotypes including root hair nuclear shape, stomatal complex development, and pollen viability. Therefore, MKAKU41 appears to act in some of the same genetic pathways as the NE-associated LINC complex proteins such as SUN and KASH. This genetic data in maize is interesting when considered with our findings that heterologous overexpression phenotypes and actin perturbations (Figs. 1–5) disrupt nuclear architecture and nuclear envelope organization. Taken together, all of these experiments establish biological roles for MKAKU41, NCH1, and NCH2 as nucleoskeletal proteins that regulate fundamental nuclear processes in cellular structure and function.

## DISCUSSION

Regulation of nucleus size and shape is important for many fundamental cellular processes in all eukaryotes. Nuclear architecture is controlled by multiple interactions involving the NE, NE-associated complexes, and the nucleoskeleton. Here, we characterized multiple maize nucleoskeletal proteins, which, like their animal counterparts, controlled nuclear dynamics. An overarching goal motivating this study is to establish the general rules that apply across the plant domain, an evolutionarily vast space. Towards this goal, we have utilized the tobacco transient heterologous expression assay as a powerful and versatile experimental platform for plant nuclear envelope research. In this study and previously, we have established cellular localization, protein-protein interactions, dose-response phenotypes, and live cell imaging that allows for kymographic analysis, mobility via FRAP, and interactions via AP-FRET, all of which have enabled and accelerated our understanding of grass and model crop NE biology (Gumber et al. 2019b, a).

The maize nucleoskeletal proteins examined in this study are NCH1, NCH2 and MKAKU41, each of which has one or more homologs (Fig. 1A) in eudicot species (Gumber et al. 2019a) and all of which exhibit nuclear localization and NE enrichment in heterologous expression systems. Interaction data from this (Fig. 3C) and prior studies further indicate that these proteins are coupled to the NE via the LINC complex. These findings, together with those from other plant species, point to the broad conservation of plant nucleoskeletal proteins across angiosperms (Goto et al. 2014; Meier et al. 2017; Ciska et al. 2019; Sakamoto 2020). The functional conservation of these components is evidenced by previously reported cross-species functional rescue (Gumber et al. 2019b) and by the current study where we show that the maize MKAKU41 (Fig. 4) interacts with Arabidopsis AtSUN2, causing altered nuclear localization of the Arabidopsis AtSUN2.

Multiple lines of evidence for conservation of plant LINC complexes are also seen at the organismal and phenotypic level. For instance, we (Fig. 7) and others found that mutant phenotypes commonly include the rounding up of root hair nuclei, disruption of stomatal complex development, and effects on pollen shape and viability (Dittmer et al. 2007; Goto et al. 2014, 2020; Zhou et al. 2014; Gumber et al. 2019b; Newman-Griffis et al. 2019). The nuclear shape defects in root hairs have become a hallmark of LINC defects in plants, and as such were predicted. However, the stomatal complex and pollen shape phenotypes have not been previously observed for plant mutants of MKAKU4 genes, but they resemble to some extent those of the plant KASH mutant, *mlks2* (Gumber et al. 2019b). To gain genetic insight into these nucleoskeletal proteins in the crop species maize, we searched for transposon-disrupted alleles of MKAKU41, NCH1, and NCH2. Of these, we found that the *MKAKU41* gene was reported to have a Mutator insertion (McCarty and Meeley 2009), described here as the first known mutant allele, *mkaku41*. The insertion site was in the 5’-UTR (Fig. 6), a common hot-spot for Mu insertion (Zhang et al. 2020). The transcript abundance was not significantly reduced in the mutants, but the mutant allele produced an extremely truncated 5’ UTR of 23 bp or less. Given that the median 5’ UTR length in maize was recently determined to be 132 bp (Leppek et al. 2018), such an extremely short 5’ UTR in the *mkaku41* mutants may abolish or greatly decrease the ability of the cell to utilize the native start codon for translation of the full-length protein. If the mutant 5’ UTR is too short for efficient ribosome assembly and scanning, the next in-frame start codon is considerably farther downstream, which would result in a loss of the first 125 AA. Additionally, the mutant 5’UTR may lack regulatory mRNA sequences in the first ~70 bases of the full-length transcript. Further genetic and experimental analyses with new alleles, gene editing, or application of specific biotic or abiotic stresses will be needed to gain a better understanding of how these plant nucleoskeletal proteins functionally interact with the genomes they help to organize.

The regulation of nuclear morphology and intra-nuclear organization was further explored to gain mechanistic insight, using the tobacco transient expression assay with fluorescently tagged proteins and quantitative microscopic analyses. We used this approach to explore multiple aspects of the remarkable nuclear architecture disruption caused by overexpression of each of the three maize nucleoskeletal proteins examined. The severity of the nuclear disruption and of the NE invaginations was increased by co-overexpression of two components (e.g. MKAKU41 with NCH1 or NCH2), or by increasing the transfecting plasmid concentration, expected to increase their expression levels. These findings (Fig. 2) and previous studies from plants and animals reveal that proper nucleoskeleton protein concentration may be a primary determinant for overall nuclear architecture (Legartová et al. 2014; Goto et al. 2014; Jorgens et al. 2017).

In addition to protein abundance, components of the nuclear invaginations were tested for the presence of LINC and ER proteins. Knowing that SUN proteins interact with ONM KASH proteins as part of the core LINC complex, we tested whether the intranuclear foci of MKAKU41-FP reflected protein aggregates of entire NE, checking for colocalization with two types of markers, those in the NE but not the LINC complex or those in the ER membrane. All of these, including multiple ONM markers, colocalized with the aberrant intranuclear structures (Figs. 4 and 5), demonstrating that these intranuclear structures contain components from both the INM and ONM of the nuclear envelope. These plant nuclei invaginations may contain, therefore, the entire NE proteome as well as NE-associated chromatin that would normally be limited to the nuclear periphery. Such invaginations are known to occur in plants and animals, which can show grooves, deformations, actin, or ER in stable structures seen as deep invaginations (Collings et al. 2000; Schermelleh et al. 2008). Interestingly, the membrane invaginations and deformations caused by MKAKU41 also resemble to some extent animal nuclear deformations associated with Lamin-A mis-expression (Lammerding et al. 2004; Schreiber and Kennedy 2013; Swift et al. 2013; Legartová et al. 2014).

In our experimental set up, we disrupted the LINC complex at two different connections to investigate the effect of these disruptions on the NE structure. The first, a SUN2ΔN, severed the LINC-to-nucleoskeleton connection; the second MLKS2ΔARM, severed the LINC-to-cytoskeleton connection. It is quite interesting that these two disruptions exhibited contrasting effects on the severity of invaginations. Our domain-deletion analyses showed that perturbation of the LINC-to-nucleoskeleton connection *reduced* the severity (Fig. 4A), whereas preventing the LINC-to-cytoskeleton connection *increased* the severity of invaginations (Fig. 5). This has important mechanobiological implications for the idea that plant nuclear shape involves a balance of forces between actin-nucleus interactions and nucleoskeletal components. Interestingly, AtKAKU4, arabidopsis KASH proteins WIPs, and NE-associated myosin are all involved in nuclear migration in various cellular processes, such as in pollen-tube growth (Goto et al., 2020, Meier et al., 2017), which also involves changes in actin dynamics.

Moving the nucleus exerts physical stress on the NE and the opposing forces of the LINC components (nucleoskeletal and cytoskeletal) are expected to be, therefore, important for maintaining NE integrity and stability (Enyedi and Niethammer 2017). The contrasting effects on NE integrity that we observe in this study are an indicator that maize nucleoskeletal components may be functionally associated with just such a tug-of-war process that manifests as regulation of nuclear shape. Multiple lines of evidence are consistent with this idea, including the change to spherical nuclei caused by genetic knockouts of a nuclear-envelope-localized myosin (Tamura et al. 2013) and the rounding up of root hair nuclei caused by the maize *mlks2* mutation (Gumber et al. 2019b). Along these lines, our study adds to the growing body of evidence that plants deploy a general mechanism for nuclear shape in which a balance of forces is achieved through LINC-interacting components on both sides of the NE, ensuring its structure and function as a flexible cellular partition. In the current study, we note multiple indications (Fig. 5) that support this tug-of-war type arrangement. These ideas align with results from mammalian studies that identify roles for the LINC complex in mediating mechanical crosstalk between the cytoplasm and nucleus (Alam et al. 2016; Jorgens et al. 2017; Hieda 2019; Agrawal and Lele 2019; Bouzid et al. 2019).

Previous investigations of CRWN and KAKU4 have focused on chromatin structure and nuclear architecture (Dittmer et al. 2007; Grob et al. 2014; Hu et al. 2019), but our studies indicate that nucleoplasmic disruptions can also affect normal developmental processes, an important finding for crop species. In animals, the interplay between cellular-level structural integrity and genomic responses to environmental and developmental processes is increasingly recognized as a complex process involving lamins as central players (Gerbino et al. 2018). This study advances our knowledge of the plant nucleoskeleton by identifying the components and their roles in regulating fundamental dynamic processes of the plant nuclear envelope.

## METHODS

### Cloning

Maize gene constructs and sequence information for the clones used in this study are listed in Table S3. NCH1 ORF was custom-synthesized with *Bam*HI and *Sbf*I at the 5’ and 3’ ends, respectively (Genscript Biotech Corporation, NJ). The *Bam*HI-NCH1-*Sbf*I construct was sub-cloned by restriction cloning into an ECGFP donor vector containing eGFP-FLAG-HA (Gumber et al. 2019, JCB), to create the eGFP-FLAG-HA-NCH1 entry vector, named NCH1ec. Similarly, the *Bam*HI-NCH2-*Pst*I construct was custom synthesized and sub-cloned into an ECGFP donor vector to create the eGFP-FLAG-HA-NCH2 entry vector, named NCH2ec. For construction of the mKAKU41vector, *Bam*HI-mCherry-FLAG-HA-MKAKU41-*Bam*H1 was synthesized by Genscript and cloned in pUC18 at the *Bam*HI restriction site. From this cloning vector, the mCherry-FLAG-HA-MKAKU41 gene construct was amplified using KAKUattF (5’-GGGGACAAGTTTGTACAAAAAAGCAGGCTTCATGGTTAGCAAGGGAGAAGAGG-3’) and KAKUattR (5’-GGGGACCACTTTGTACAAGAAAGCTGGGTCTCACGTAGCCCGTCCCCGT-3’) primers and inserted into pDONR221 vector by BP cloning (Invitrogen), to generate the MKAKU41 entry clone, named MKAKU41ec. For the generation of plant expression vectors, the fluorescent fusion protein constructs from these three entry clones were then transferred individually to the destination vector pH7WG2 (Karimi et al. 2002) by Gate LR recombination (Invitrogen).

For production of the p35S∷SP-mCherry-GFP-HDEL positive control for apFRET, mCherry was PCR amplified with Q5 polymerase (NEB) using primers JM403 (5’-GGGGACAAGTTTGTACAAAAAAGCAGGCTACAATGAAAGCCTTCACACTCGCTCTC TTCTTAGCTCTTTCCCTCTATCTCCTGCCCAATCCAGCCATGGTGAGCAAGGGCGA GGAGG-3’) & JM404 (5’-acctccactgccaccCTTGTACAGCTCGTCCATGCCG-3’). The primers included both the Gateway cloning attB1 site and secretion signal at the 5’ end, and a 15nt overhang for Gibson assembly at the 3’ end, which contained a GGSGG amino acid linker between mCherry and GFP upon fusion. GFP was amplified using primers JM367 (5’GGGGACCACTTTGTACAAGAAAGCTGGGTGtcataattcatcatGCTTGTACAGCTCGTCCATGCCGAGAG-3’) & JM405 (5’GTACAAGggtggcagtggaggtATGGTGAGCAAGGGCGAGGAGC-3’) which contained the GGSGG linker at the 5’ end, and the HDEL ER retention motif, an attB2 site, and a stop codon at the 3’ end. These two products were fused together using the NEB HIFI Gibson assembly enzyme mix and incubated at 50°C for one hour. A Gateway BP reaction into pDONR221 and subsequent LR reaction into pB7FWG2 was then performed to produce the final vector. All steps confirmed by colony PCR and sequencing.

### Agrobacterium transformation

Constructs were transformed into A. tumefaciens GV3101. Transformation was performed by incubating plasmid DNA and chemically competent agrobacterium on ice for 30 minutes, followed by 5 minutes cold shock in liquid nitrogen, then 5 minutes heat shock at 37□ in a rotating incubator. After heat shock, 200μL LB media was added and cells incubated at 28□ for two hours. Cells were then plated in LB plates containing Spectinomycin (50μg/mL), Gentamycin (10 μg/mL) and Rifampicin (25 μg/mL) and incubated for two days at 28°C. Individual colonies were then picked, grown O/N and transformed into *N. benthamiana*.

### Plant growth conditions

*N. benthamiana* plants were grown in 16:8h light:dark cycle in a greenhouse maintained at 21 °C. Infiltrated plants were 5-6 weeks old.

### Live cell imaging

Fluorescently tagged proteins of interest were transiently transformed into *N. benthamiana* as described previously (Sparkes et al. 2006). Protein expression constructs first reported here are p35S∷NCH1-GFP, p35S∷NCH2-GFP and p35S∷MKAKU41-mCherry. All other markers have been previously published: p35S∷GFP-CNX (Irons et al. 2003), p35S∷LBR-GFP (Irons et al., 2003), p35S∷MLKP1-GFP (Gumber et al. 2019a), p35S∷YFP-AtSUN2 (Graumann et al. 2010), p35S∷YFP-AtSUN2ΔNterm (Graumann et al. 2010), p35S∷MLKS2-GFP and p35S∷MLKS2ΔARM-GFP (Gumber et al. 2019b). An agrobacterium culture of OD 0.1 was used in all conditions unless otherwise stated and cells were imaged three days after transformation. The GFP / mCherry combinations were imaged using a Zeiss LSM 800 confocal microscope with line switching, 488nm and 561 nm excitation, and 500-550nm and 565-620nm emissions, collected for GFP and mCherry respectively. For GFP / YFP and YFP / mCherry imaging, a Zeiss LSM 880 was used with frame switching. For GFP / YFP imaging, 488nm and 514nm excitation was used with emission collected between 500-550 and 525-560 for GFP and YFP respectively. For YFP / mCherry imaging, 514nm and 561nm excitation was used, and emission collected between 517-560nm and 561-624 nm respectively. An image size of 512×512 pixels with a scan zoom of 4 and a 63x 1.4NA lens was used for all imaging described above. All combinations were performed with three independent experimental repeats; representative images are shown. For Fluorescence recovery after photobleaching (FRAP) a 100x 1.4NA lens was used with a 4μm ROI in the center of the image, encompassing the nucleus. Five scans were taken pre-bleach and then the 488nm laser bleached the ROI by using 100% transmission for 20 iterations. Recovery Images were then collected for one minute to monitor recovery. Data was normalised and FRAP curves produced as described previously (Martiniere et al. 2012).

For acceptor photobleaching förster resonance energy transfer (apFRET) a 100x 1.4NA lens was used with a 4μm ROI in the center of the image, encompassing the nucleus. Five scans were performed with both GFP and mCherry emission / excitation, and then the mCherry construct was bleached in the ROI by the 561nm laser at 100% transmission for 20 iterations. Following this, five post-bleach scans were taken. Data was normalized and apFRET efficiency (%) calculated as previously (Graumann et al. 2010; Graumann 2014; Pawar et al. 2016). A minimum of 30 nuclei per condition were used for apFRET across three experimental repeats. A one-way ANOVA was performed to determine statistically significant differences between samples. For Latrunculin-B (LatB) treatment for depolymerisation of the actin cytoskeleton, samples were incubated with 25 μM LatB for one hour, as this has previously been shown to depolymerise the actin cytoskeleton sufficiently (McKenna et al. 2019). Graphs were generated with graphpad and as described in the figure legends.

### Maize plant material and genotyping

The wild-type W22 used in this study is a color-converted W22 line obtained from Hugo Dooner (Waksman Inst., Rutgers, New Jersey, USA) derived by (Brink 1956). The UF-Mu-00395 seed stock was obtained from the Maize Genetics Cooperation Stock Center (http://maizecoop.cropsci.uiuc.edu/). The plants were grown at the Florida State University Mission Road Research Facility (Tallahassee, FL, USA) during summer 2017 and 2018, and propagated by out-crossing to W22. In the fall of 2018, the progeny seeds were grown in the greenhouse in the King Life Sciences Building (Biological Science Dept, Florida State University, Tallahassee, FL, USA). The segregating plants were self-crossed to obtain mutant plants from among the progeny.

DNA was isolated from 4-week old seedlings as described previously in (Gumber et al. 2019b). PCR genotyping was carried out using a combination of gene-specific forward (F, 5’-CCCGTGAAGCCGAAGGCAGA-3’) and reverse (R, 5’-CGCCTCACGCTCACGCTCAC-3’) primers, or transposon-specific Tir6 primer (5’-GAGAAGCCAACGCCAWCGCCTCYATTTCGTC-3’) in combination with F or R primer. The PCR products were resolved by agarose gel electrophoresis and cloned in pCRTM4Blunt-TOPO® Vector (Invitrogen cat # K2875-20) by TA cloning. The clones were sequenced and the insert sequences were verified using M13F and M13R vector primers at the Molecular Cloning Facility, Department of Biological Sciences, Florida State University. The sequences were aligned with the W22v2 reference genome to validate the transposon insertion site.

### Microscopy in maize

Maize root hair imaging was carried out as described in (Gumber et al. 2019b). Briefly, roots were harvested from 5-day old seedlings and fixed for 1 hour in Buffer A (Howe et al. 2013) supplemented with 4% paraformaldehyde. Small sections of root tissue containing root hair were stained with 3 μg/mL DAPI for 20 min at room temperature, mounted with VECTASHIELD, and imaged on an EVOS fluorescence microscope (Thermo Fisher Scientific). The images were processed using the Analyze Particle function of ImageJ to measure the longest diameter and circularity of the nuclei.

For stomatal complex imaging, plants were grown in the greenhouse and the 4th leaf was harvested at its first appearance. The harvested leaf was fixed in Buffer A with 4% paraformaldehyde for an hour at room temperature with rotation. The tissue was rinsed thrice with and stored in Buffer A at 4C, until further use. The leaf tissue was placed on a glass slide, chopped into small pieces and stained with 3 μg/mL DAPI for 20 min at room temperature, mounted with VECTASHIELD, and imaged on an EVOS fluorescence microscope (Thermo Fisher Scientific).

Pollen grain staining was carried out as previously described in (Gumber et al. 2019b). Briefly, male flowers were harvested before dehiscence and fixed in Carnoy’s fixative (6 alcohol:3 chloroform:1 acetic acid) for a minimum of 2 hours at room temperature. Anthers were extruded from flowers with the help of a micro scalpel and forceps on a glass slide. Staining was carried out with modified Alexander’s stain containing Malachite green (0.01%), Acid Fuchsin (0.05%) and Orange G (0.005%) as described to differentiate viable (magenta) pollen grains from aborted (green) pollen grains. Bright field images of the pollen grains were collected on Revolve microscope (Echo Labs). At least 300 pollen grains each from 3 plants of every genotype were counted to calculate pollen viability. Pollen roundness was carried out using Fiji.

### RNA isolation and library preparation

Segregating wildtype and mutant mkaku41 plants were grown in the greenhouse. From two week-old plants, fourth leaves were harvested and from 6-8 week old plants, mid-prophase meiotic-staged male flowers were harvested. The tissues were immediately stored in liquid nitrogen. RNA was isolated from three biological replicates for each genotype using Qiagen RNeasy Plant mini kit per manufacturer’s instructions. Integrity of the RNA was tested using the Bioanalyzer (Agilent) system. For library preparation, sample input was 400 ng total RNA (determined by Qubit RNA HS reagents, Thermo) with RIN >7 (Bioanalyzer RNA Nano, Agilent). Libraries were prepared with the Biomek 400 Automated Workstation (Beckman Coulter), using the NEBNEXT Ultra II RNA Library Prep kit for Illumina (New England Biolabs) according to manufacturer’s instructions, with an RNA fragmentation time of 15 minutes, a 1/10^th^ dilution of NEB adaptor and 11 cycles of PCR amplification with dual-indexing primers. Amplified libraries were initially quantified by Qubit DNA HS reagents, checked for size and artifacts using Bioanalyzer DNA HS reagents, and KAPA qPCR (KAPA Biosystems) was used to determine molar quantities of each library. Individual libraries were diluted and pooled equimolar, and the pool was again checked by Bioanalyzer and KAPA qPCR before submission for sequencing.

### RNA sequencing and data analysis

RNA-seq libraries were sequenced on a Novaseq 6000 at the Translational Science Lab, College of Medicine, Florida State University. Approximately 40 million single-end 100 base reads were obtained for each biological replicate in this experiment and are available from NCBI sequence read archive project, accession number PRJNA675860. Contaminating 3’ adapter sequences were trimmed from the demultiplexed raw reads using cutadapt version 1.16. Raw and trimmed reads were subjected to quality control testing with fastqc. Trimmed reads were aligned to the W22 genome assembly “Zm-W22-REFERENCE-NRGENE-2.0” using the splice-aware aligner hisat2. Briefly, Hisat2 indices were constructed from known exons, and splice sites extracted from the W22 genome annotation (Zm00004b) and the reference genome assembly (Zm-W22-REFERENCE-NRGENE-2.0.fasta). Trimmed reads were then aligned to the resulting splice-aware hisat2 index using the following optional arguments: --rna-strandness R, --dta-cufflinks, --summary-file. Predicted novel transcripts were assembled and merged across replicates and samples using stringtie2 in “conservative” mode. Per-transcript coverage tables were prepared by stringtie2 in “ballgown” format. Resulting coverage tables were converted into count tables suitable for differential expression analysis by DEseq2 in R using the tximport package. Differential expression analysis was performed separately for each tissue group i.e (leaf mutant vs. WT and tassel mutant vs. WT). Briefly, genes with fewer than 10 counts across all replicates were discarded and DEseq2 results were generated for both tissue groups such that log2(fold-change) estimates were reported for (mutant/WT) ratios. Statistically significant differentially expressed (adjusted p-value <0.05) genes were subsequently extracted from each of the resulting DEseq2 tables for further analysis (Table S2).

## Acknowledgments

We would like to acknowledge the Bioimaging unit at Oxford Brookes for access to the confocal microscopes. This work was performed as part of the Cost action # CA16212 ‘INDEPTH’ whose members are appreciated for their fruitful discussions. This work was supported in part by a grant to HWB from the National Science Foundation (NSF IOS 1444532).

## Legends for Supplemental Figures

**Figure S1:**
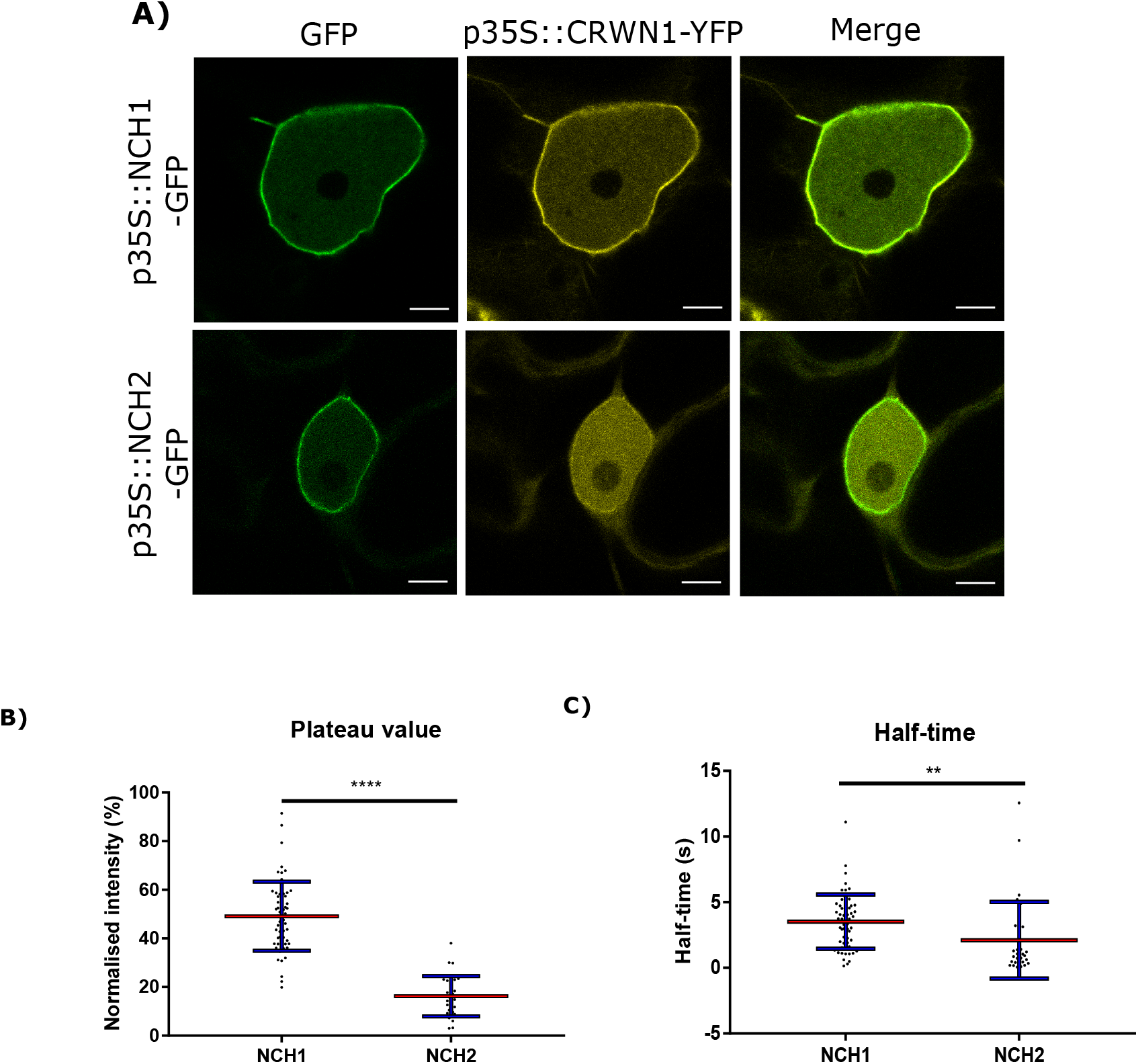
The Maize NCH1 and NCH2 proteins co-localize with their Arabidopsis homolog CRWN1. Confocal live cell imaging of cells co-expressing either NCH1 or NCH2 with Arabidopsis CRWN1 shows colocalization. B) Scatter dot plot of plateau recovery values from NCH1 and NCH2 FRAP time course experiment. C) Scatter dot plot of halftime recovery values from NCH1 and NCH2 FRAP time course experiment. Scatter dot plot of NCH2 Scale bar denotes 5μm. N≥ 30 nuclei imaged across three experimental replicates.

**Figure S2:**
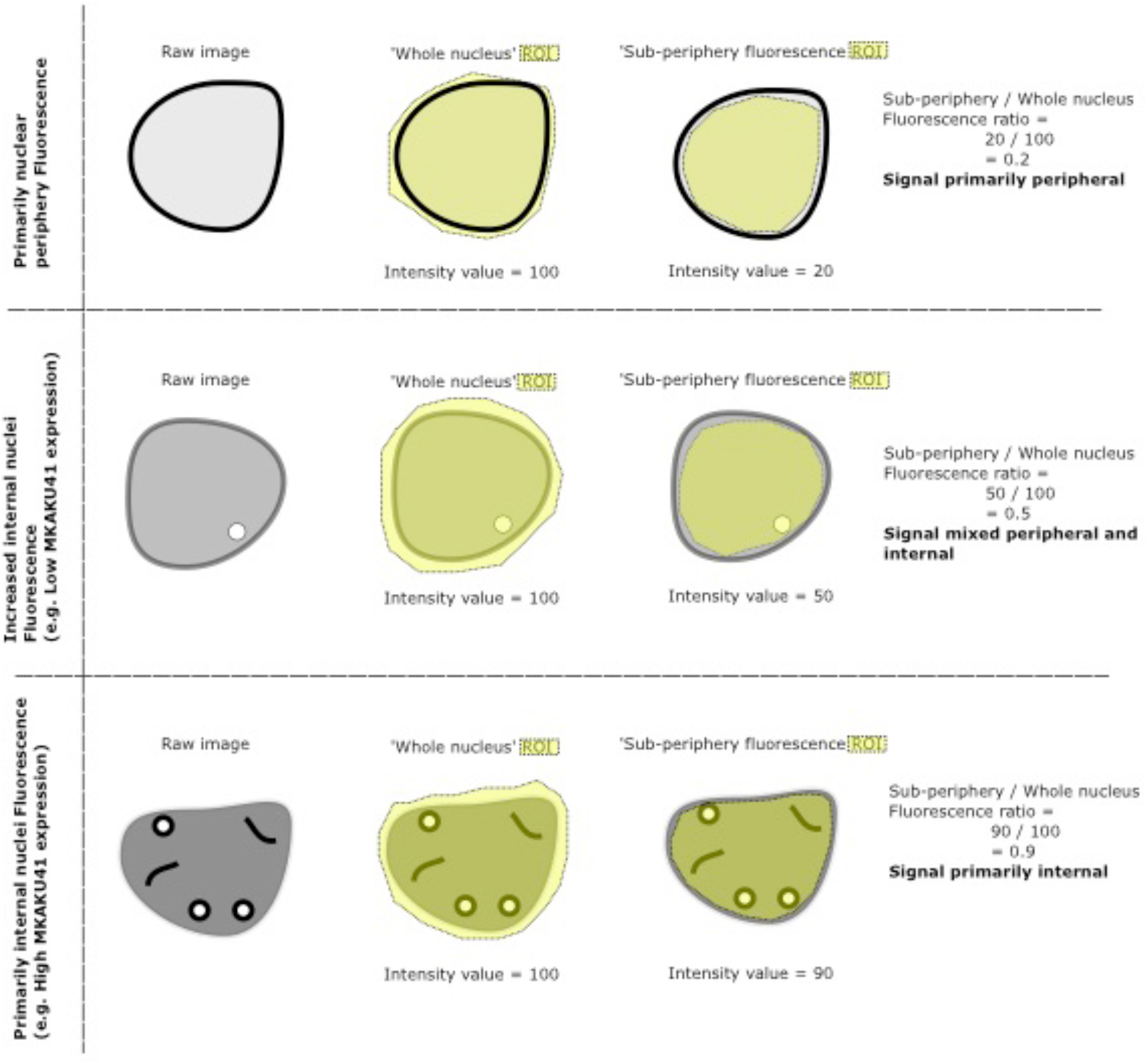
Description of Sub-periphery / whole nuclei fluorescence ratio measurement. Diagram describing how the sub-periphery over whole nuclei measurement was determined using image J. Two regions of interest (ROIs) were generated, one encompassing the whole nuclei and one only sub-periphery nuclear fluorescence (yellow ROIs with hashed boundaries). The sub-periphery fluorescence value was then divided by the whole nuclei fluorescence value in order to obtain the ratio. If the majority of fluorescence is located at the periphery / nuclear envelope, this would result in a low ratio, conversely if most fluorescence was internal, this would result in a higher ratio.

**Figure S3:**
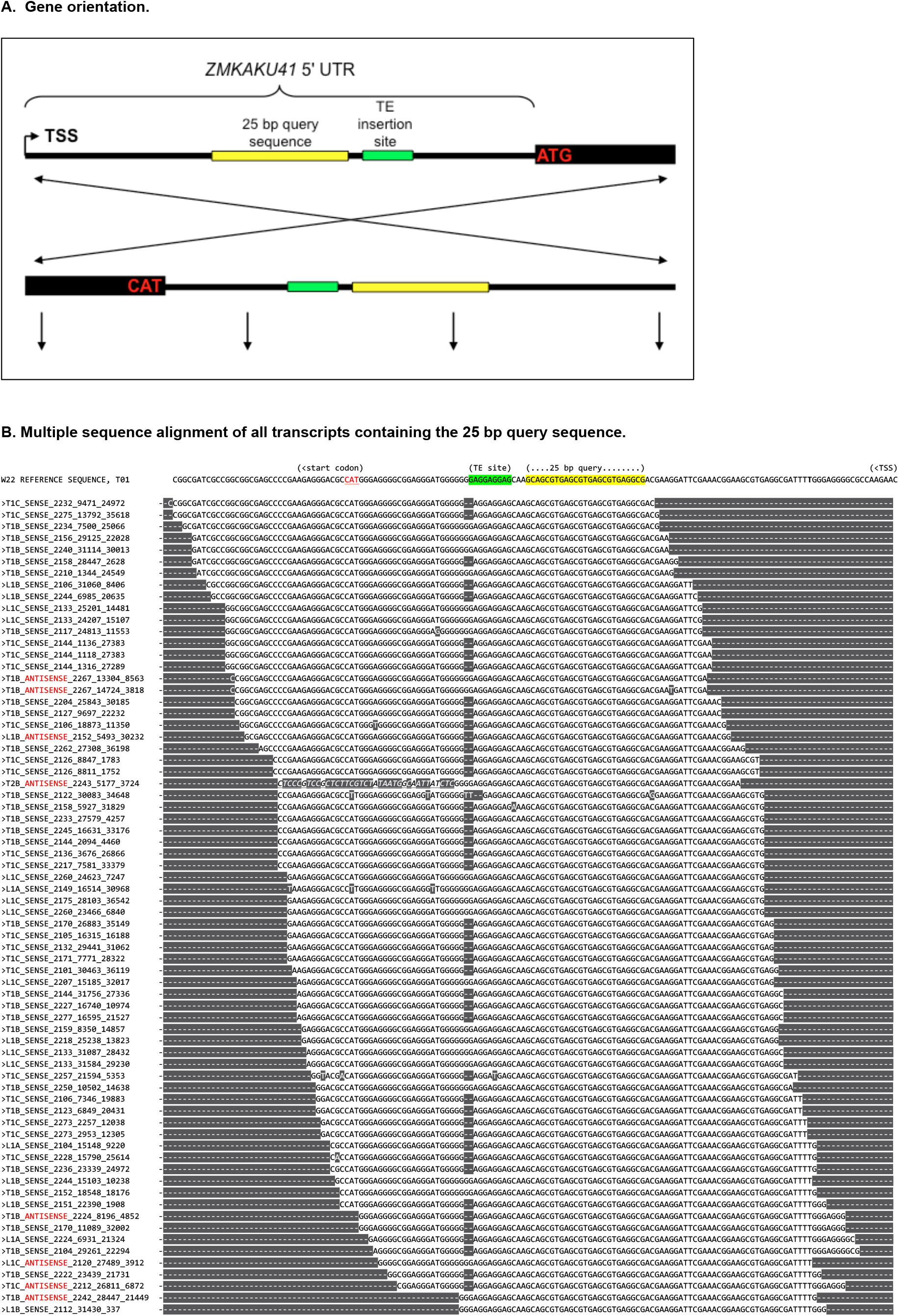
Multiple Seq Alignment of Transcripts. A) The 5’ UTR region and a small portion of the CDS are diagrammed as shown in Figure 5, and reversed (bottom configuration) as aligned in the multiple sequence alignment. B) The multiple sequence alignment displays all of the RNA-seq reads with a perfect match to the 25 bp query sequence (yellow) using grep of the fastq files. All matches were converted to FASTA sequences for multiple sequence alignment. The reference genome sequence is shown at top for comparison. The sequence identifiers start with single characters for tissue (“L” for leaf; “T” for tassel), genotype (“1” for wildtype, “2” for mkaku41 homozygous mutant), or bioreplicate (“A”, “B”, or “C” for bioreplicate 1, 2, or 3, respectively), followed by unique identifier from Illumina sequence read name. The strandedness is indicated relative to the gene model, with all antisense RNAs indicated (ANTISENSE, red text).

## Supplementary Tables

**Table S1:** Values for apFRET efficiency with Internal controls and multiple comparisons of all treatments. A) All apFRET efficiency % values for data presented in figure 3C including internal control values. B) Tukey multiple comparison one way Anova dataset from all apFRET data presented, including that presented in figure 3C.

| A) | NCH1 & CNX | NCH1 & CNX IC | NCH1 & KAKU41 | NCH1 & KAKU41 IC | NCH2 & CNX | NCH2 & CNX IC | NCH2 & KAKU41 | NCH2 & KAKU41 IC | CNX & KAKU41 | CNX & KAKU41 IC | mCherry -GFP -HDEL | mCherry -GFP -HDEL IC |
| --- | --- | --- | --- | --- | --- | --- | --- | --- | --- | --- | --- | --- |
| Number of values | 30 | 30 | 30 | 30 | 30 | 30 | 33 | 33 | 30 | 30 | 30 | 30 |
| Mean FRET Efficiency (%) | 1.4 | -0.67 | 10 | -0.94 | 3.3 | -0.86 | 11 | -0.76 | 6.6 | -2.1 | 19 | -0.9 |
| Std. Deviation | 3.7 | 0.53 | 2.4 | 0.3 | 2.7 | 0.4 | 3 | 0.35 | 4.6 | 0.63 | 6.8 | 1.3 |
| Std. Error of Mean | 0.67 | 0.097 | 0.43 | 0.055 | 0.5 | 0.074 | 0.53 | 0.061 | 0.84 | 0.11 | 1.2 | 0.23 |

| Tukey's multiple comparisons test | Mean Diff. | 95.00% CI of diff. | Below thr | Summary | pd P Value |
| --- | --- | --- | --- | --- | --- |
| NCH1 & CNX vs. NCH1 & CNX IC | 2.073 | -0.4343 to 4.580 | No | ns | 0.2208 |
| NCH1 & CNX vs. NCH1 & KAKU41 | -8.939 | -11.45 to -6.432 | Yes | **** | <0.0001 |
| NCH1 & CNX vs. NCH1 & KAKU41 IC | 2.343 | -0.1640 to 4.850 | No | ns | 0.0925 |
| NCH1 & CNX vs. NCH2 & CNX | -1.874 | -4.381 to 0.6331 | No | ns | 0.3694 |
| NCH1 & CNX vs. NCH2 & CNX IC | 2.263 | -0.2439 to 4.770 | No | ns | 0.122 |
| NCH1 & CNX vs. NCH2 & KAKU41 | -9.607 | -12.06 to -7.158 | Yes | **** | <0.0001 |
| NCH1 & CNX vs. NCH2 & KAKU41 IC | 2.163 | -0.2862 to 4.612 | No | ns | 0.1434 |
| NCH1 & CNX vs. CNX & KAKU41 | -5.199 | -7.706 to -2.692 | Yes | **** | <0.0001 |
| NCH1 & CNX vs. CNX & KAKU41 IC | 3.473 | 0.9662 to 5.980 | Yes | *** | 0.0004 |
| NCH1 & CNX vs. mCherry-GFP -HDEL | -17.14 | -19.65 to -14.64 | Yes | **** | <0.0001 |
| NCH1 & CNX vs. mCherry-GFP -HDEL IC | 2.303 | -0.2041 to 4.810 | No | ns | 0.1065 |
| NCH1 & CNX IC vs. NCH1 & KAKU41 | -11.01 | -13.52 to -8.505 | Yes | **** | <0.0001 |
| NCH1 & CNX IC vs. NCH1 & KAKU41 IC | 0.2703 | -2.237 to 2.777 | No | ns | >0.9999 |
| NCH1 & CNX IC vs. NCH2 & CNX | -3.946 | -6.453 to -1.440 | Yes | **** | <0.0001 |
| NCH1 & CNX IC vs. NCH2 & CNX IC | 0.1905 | -2.316 to 2.697 | No | ns | >0.9999 |
| NCH1 & CNX IC vs. NCH2 & KAKU41 | -11.68 | -14.13 to -9.230 | Yes | **** | <0.0001 |
| NCH1 & CNX IC vs. NCH2 & KAKU41 IC | 0.09053 | -2.359 to 2.540 | No | ns | >0.9999 |
| NCH1 & CNX IC vs. CNX & KAKU41 | -7.272 | -9.779 to -4.765 | Yes | **** | <0.0001 |
| NCH1 & CNX IC vs. CNX & KAKU41 IC | 1.401 | -1.106 to 3.907 | No | ns | 0.7958 |
| NCH1 & CNX IC vs. mCherry-GFP -HDEL | -19.22 | -21.72 to -16.71 | Yes | **** | <0.0001 |
| NCH1 & CNX IC vs. mCherry-GFP -HDEL IC | 0.2303 | -2.277 to 2.737 | No | ns | >0.9999 |
| NCH1 & KAKU41 vs. NCH1 & KAKU41 IC | 11.28 | 8.775 to 13.79 | Yes | **** | <0.0001 |
| NCH1 & KAKU41 vs. NCH2 & CNX | 7.065 | 4.558 to 9.572 | Yes | **** | <0.0001 |
| NCH1 & KAKU41 vs. NCH2 & CNX IC | 11.2 | 8.695 to 13.71 | Yes | **** | <0.0001 |
| NCH1 & KAKU41 vs. NCH2 & KAKU41 | -0.6675 | -3.117 to 1.782 | No | ns | 0.9991 |
| NCH1 & KAKU41 vs. NCH2 & KAKU41 IC | 11.1 | 8.653 to 13.55 | Yes | **** | <0.0001 |
| NCH1 & KAKU41 vs. CNX & KAKU41 | 3.74 | 1.233 to 6.247 | Yes | **** | <0.0001 |
| NCH1 & KAKU41 vs. CNX & KAKU41 IC | 12.41 | 9.905 to 14.92 | Yes | **** | <0.0001 |
| NCH1 & KAKU41 vs. mCherry-GFP -HDEL | -8.204 | -10.71 to -5.698 | Yes | **** | <0.0001 |
| NCH1 & KAKU41 vs. mCherry-GFP -HDEL IC | 11.24 | 8.735 to 13.75 | Yes | **** | <0.0001 |
| NCH1 & KAKU41 IC vs. NCH2 & CNX | -4.217 | -6.724 to -1.710 | Yes | **** | <0.0001 |
| NCH1 & KAKU41 IC vs. NCH2 & CNX IC | -0.07984 | -2.587 to 2.427 | No | ns | >0.9999 |
| NCH1 & KAKU41 IC vs. NCH2 & KAKU41 | -11.95 | -14.40 to -9.500 | Yes | **** | <0.0001 |
| NCH1 & KAKU41 IC vs. NCH2 & KAKU41 IC | -0.1798 | -2.629 to 2.270 | No | ns | >0.9999 |
| NCH1 & KAKU41 IC vs. CNX & KAKU41 | -7.542 | -10.05 to -5.035 | Yes | **** | <0.0001 |
| NCH1 & KAKU41 IC vs. CNX & KAKU41 IC | 1.13 | -1.377 to 3.637 | No | ns | 0.9444 |
| NCH1 & KAKU41 IC vs. mCherry-GFP -HDEL | -19.49 | -21.99 to -16.98 | Yes | **** | <0.0001 |
| NCH1 & KAKU41 IC vs. mCherry-GFP -HDEL IC | -0.04004 | -2.547 to 2.467 | No | ns | >0.9999 |
| NCH2 & CNX vs. NCH2 & CNX IC | 4.137 | 1.630 to 6.644 | Yes | **** | <0.0001 |
| NCH2 & CNX vs. NCH2 & KAKU41 | -7.733 | -10.18 to -5.284 | Yes | **** | <0.0001 |
| NCH2 & CNX vs. NCH2 & KAKU41 IC | 4.037 | 1.588 to 6.486 | Yes | **** | <0.0001 |
| NCH2 & CNX vs. CNX & KAKU41 | -3.325 | -5.832 to -0.8183 | Yes | ** | 0.001 |
| NCH2 & CNX vs. CNX & KAKU41 IC | 5.347 | 2.840 to 7.854 | Yes | **** | <0.0001 |
| NCH2 & CNX vs. mCherry-GFP -HDEL | -15.27 | -17.78 to -12.76 | Yes | **** | <0.0001 |
| NCH2 & CNX vs. mCherry-GFP -HDEL IC | 4.177 | 1.670 to 6.684 | Yes | **** | <0.0001 |
| NCH2 & CNX IC vs. NCH2 & KAKU41 | -11.87 | -14.32 to -9.421 | Yes | **** | <0.0001 |
| NCH2 & CNX IC vs. NCH2 & KAKU41 IC | -0.09992 | -2.549 to 2.349 | No | ns | >0.9999 |
| NCH2 & CNX IC vs. CNX & KAKU41 | -7.462 | -9.969 to -4.955 | Yes | **** | <0.0001 |
| NCH2 & CNX IC vs. CNX & KAKU41 IC | 1.21 | -1.297 to 3.717 | No | ns | 0.9125 |
| NCH2 & CNX IC vs. mCherry-GFP -HDEL | -19.41 | -21.91 to -16.90 | Yes | **** | <0.0001 |
| NCH2 & CNX IC vs. mCherry-GFP -HDEL IC | 0.0398 | -2.467 to 2.547 | No | ns | >0.9999 |
| NCH2 & KAKU41 vs. NCH2 & KAKU41 IC | 11.77 | 9.380 to 14.16 | Yes | **** | <0.0001 |
| NCH2 & KAKU41 vs. CNX & KAKU41 | 4.408 | 1.958 to 6.857 | Yes | **** | <0.0001 |
| NCH2 & KAKU41 vs. CNX & KAKU41 IC | 13.08 | 10.63 to 15.53 | Yes | **** | <0.0001 |
| NCH2 & KAKU41 vs. mCherry-GFP -HDEL | -7.537 | -9.986 to -5.088 | Yes | **** | <0.0001 |
| NCH2 & KAKU41 vs. mCherry-GFP -HDEL IC | 11.91 | 9.460 to 14.36 | Yes | **** | <0.0001 |
| NCH2 & KAKU41 IC vs. CNX & KAKU41 | -7.362 | -9.812 to -4.913 | Yes | **** | <0.0001 |
| NCH2 & KAKU41 IC vs. CNX & KAKU41 IC | 1.31 | -1.139 to 3.759 | No | ns | 0.8385 |
| NCH2 & KAKU41 IC vs. mCherry-GFP -HDEL | -19.31 | -21.76 to -16.86 | Yes | **** | <0.0001 |
| NCH2 & KAKU41 IC vs. mCherry-GFP -HDEL IC | 0.1397 | -2.310 to 2.589 | No | ns | >0.9999 |
| CNX & KAKU41 vs. CNX & KAKU41 IC | 8.672 | 6.165 to 11.18 | Yes | **** | <0.0001 |
| CNX & KAKU41 vs. mCherry-GFP -HDEL | -11.94 | -14.45 to -9.438 | Yes | **** | <0.0001 |
| CNX & KAKU41 vs. mCherry-GFP -HDEL IC | 7.502 | 4.995 to 10.01 | Yes | **** | <0.0001 |
| CNX & KAKU41 IC vs. mCherry-GFP -HDEL | -20.62 | -23.12 to -18.11 | Yes | **** | <0.0001 |
| CNX & KAKU41 IC vs. mCherry-GFP -HDEL IC | -1.17 | -3.677 to 1.337 | No | ns | 0.9296 |
| mCherry-GFP -HDEL vs. mCherry-GFP -HDEL IC | 19.45 | 16.94 to 21.95 | Yes | **** | <0.0001 |

\* Note IC = Internal FRET efficiency control
Supplementary Table S1 is from McKenna, Gumber, et al., "Maize (Zea mays L.) nucleoskeletal proteins regulate nuclear envelope remodeling...", submitted 12/2020

**Table S2:**
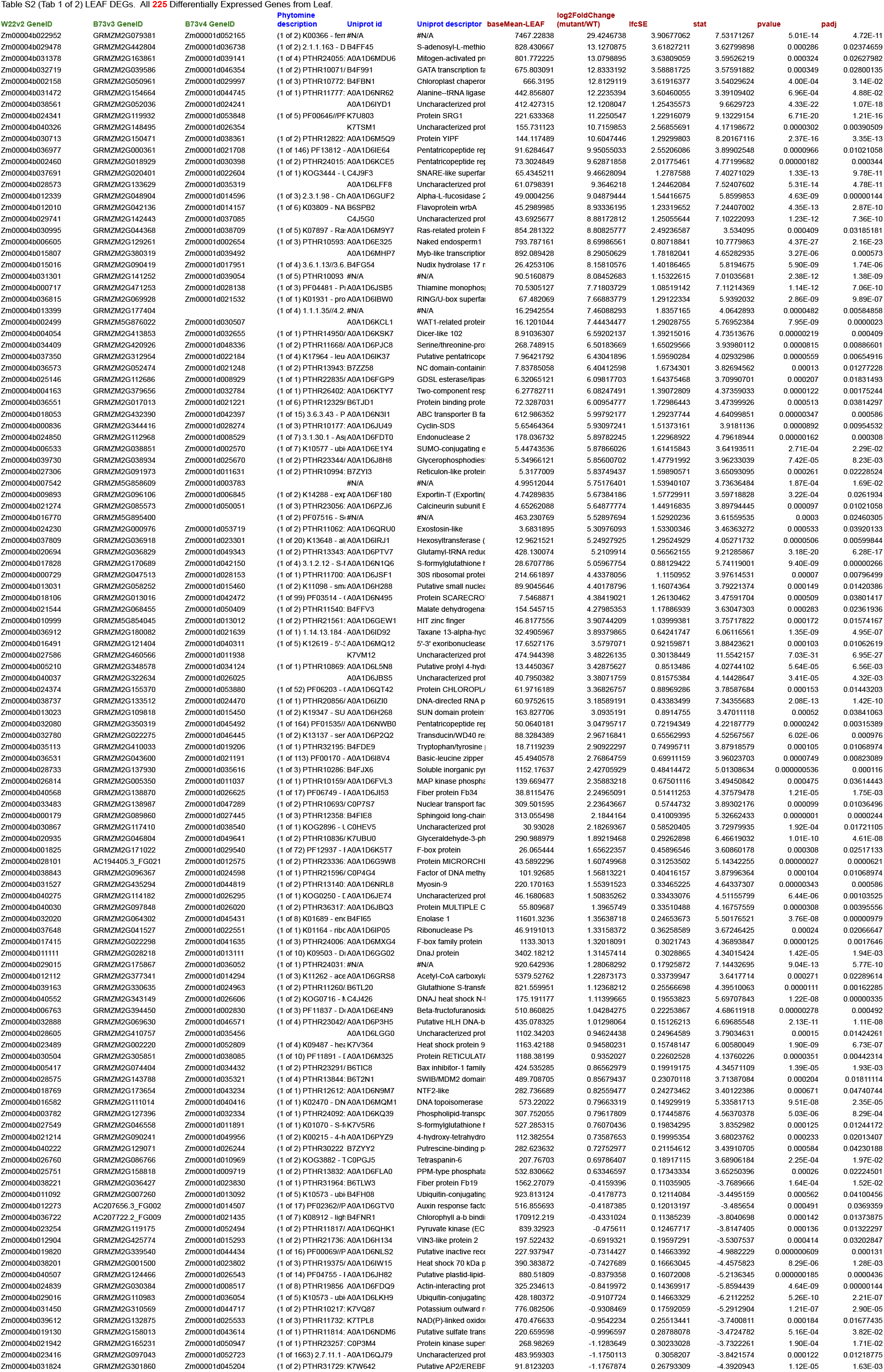
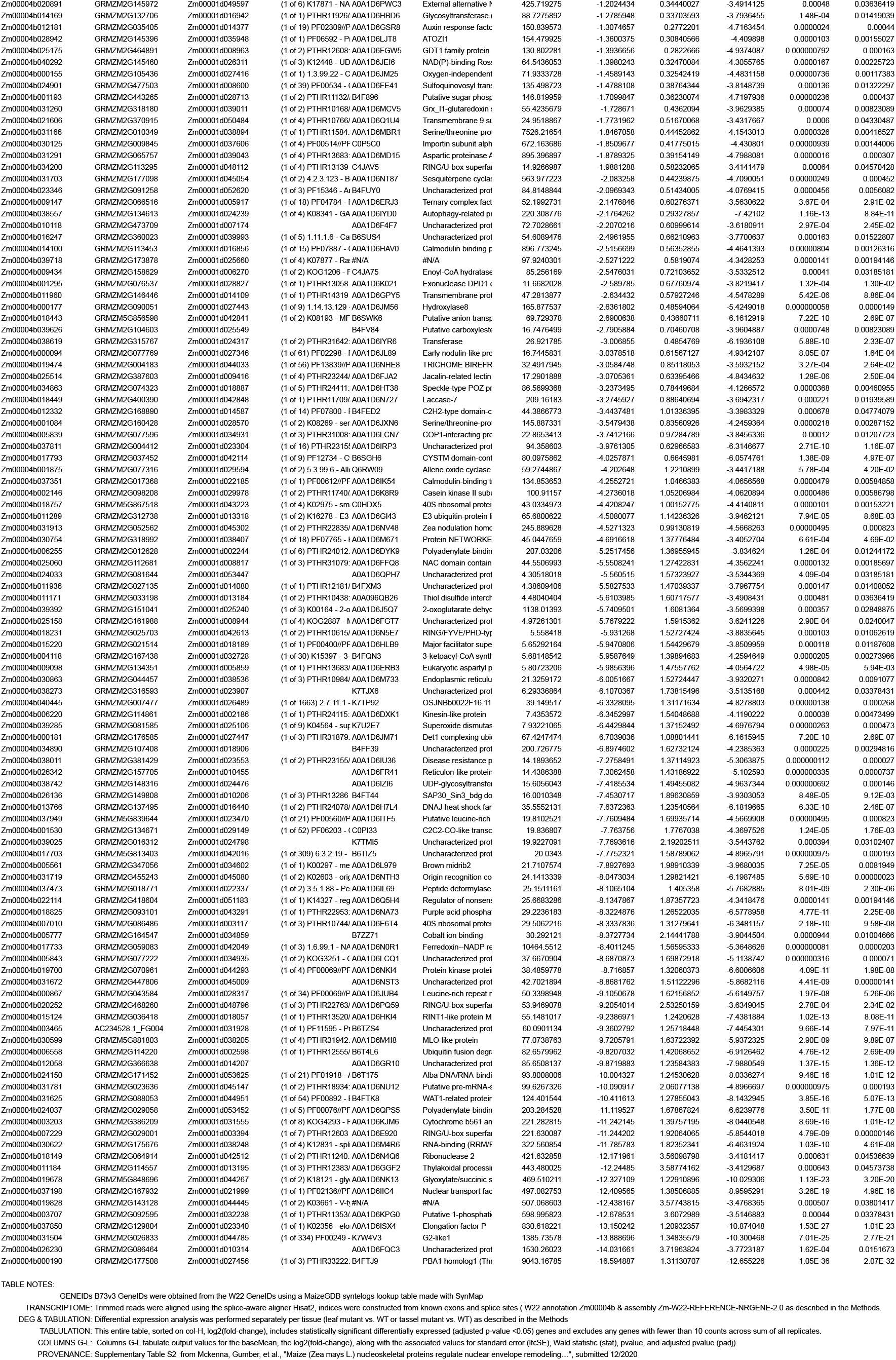
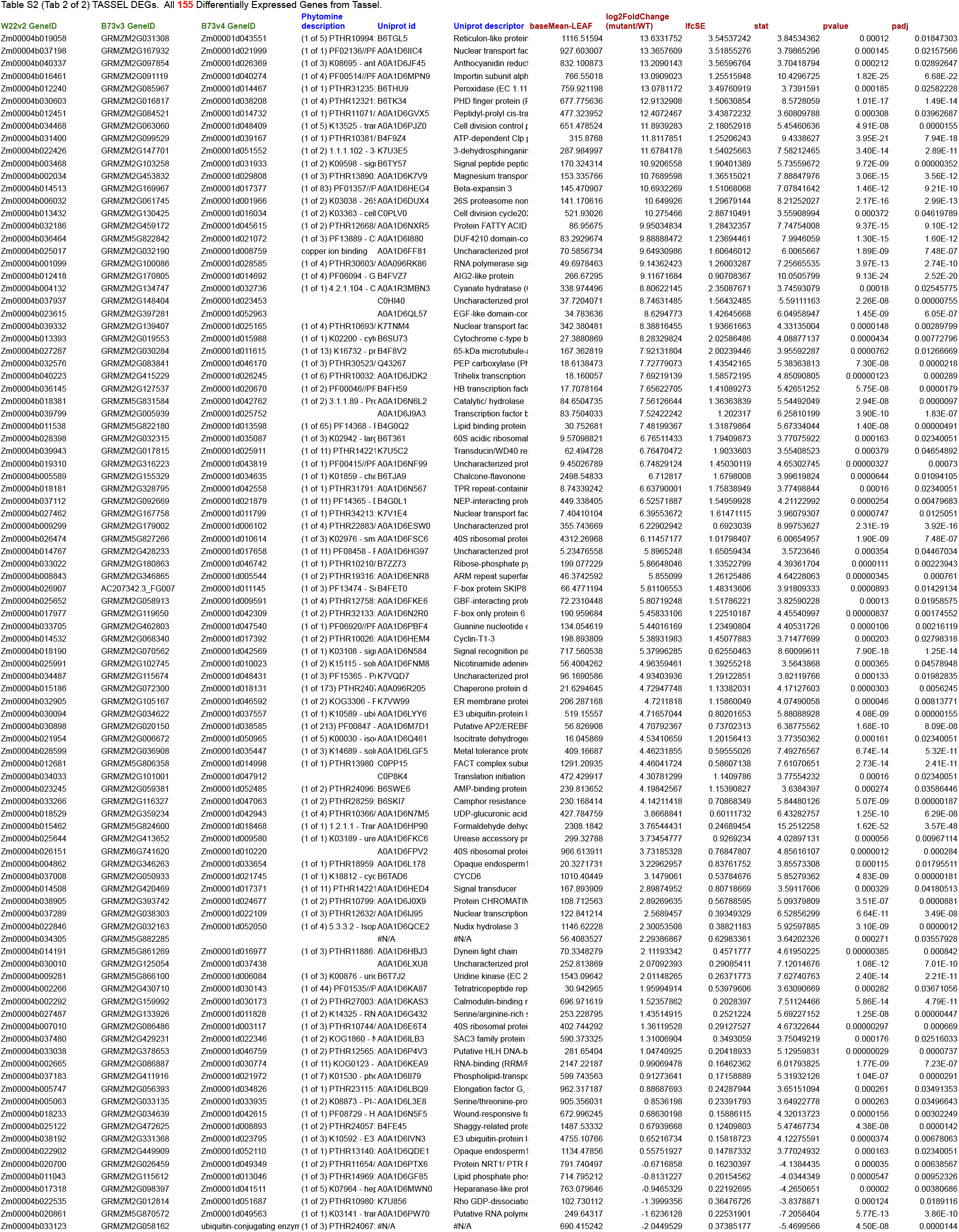
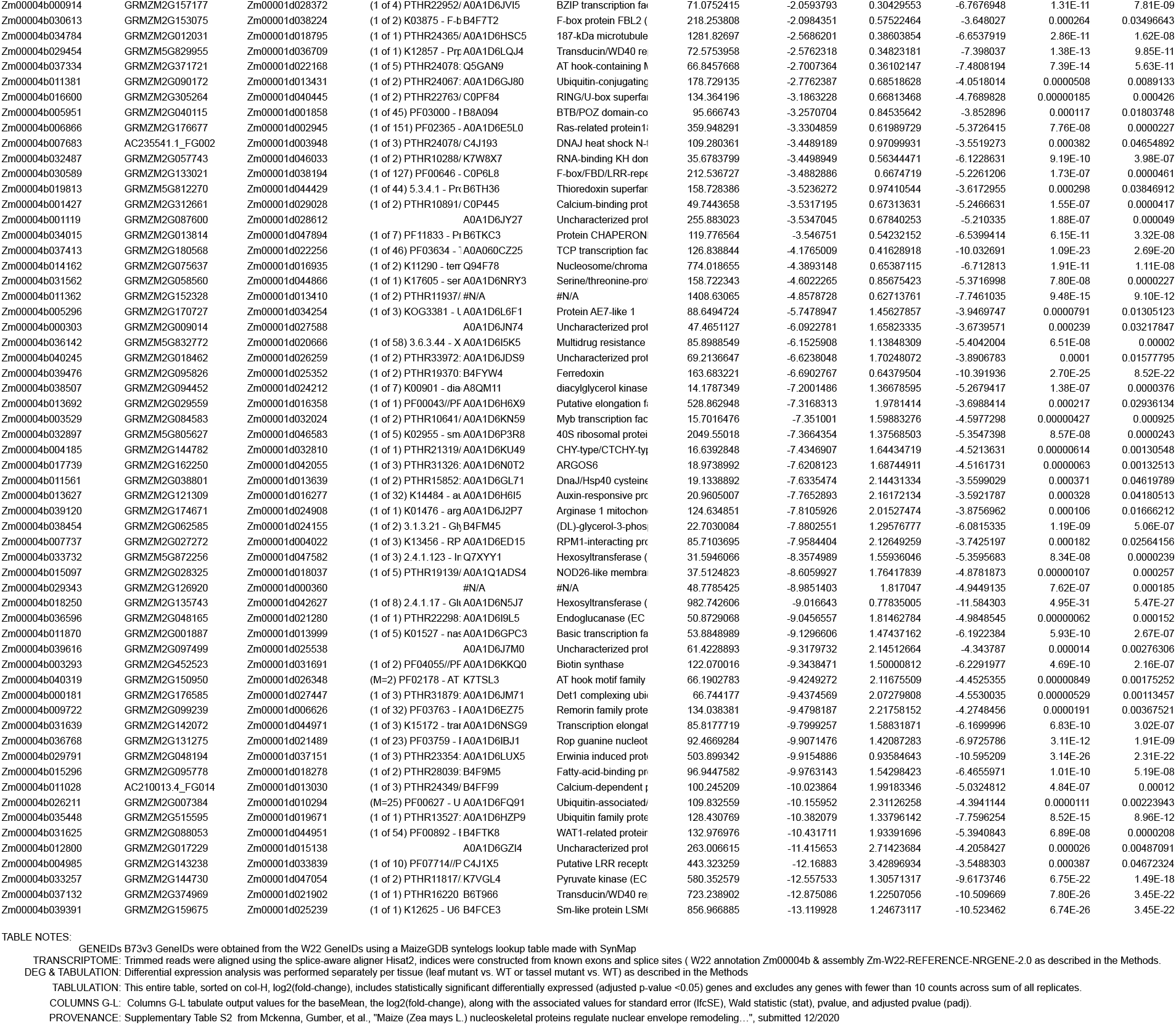
Differentially expressed genes between mkaku41 mutant versus wildtype plants for maize leaf and tassel.

**Table S3:**
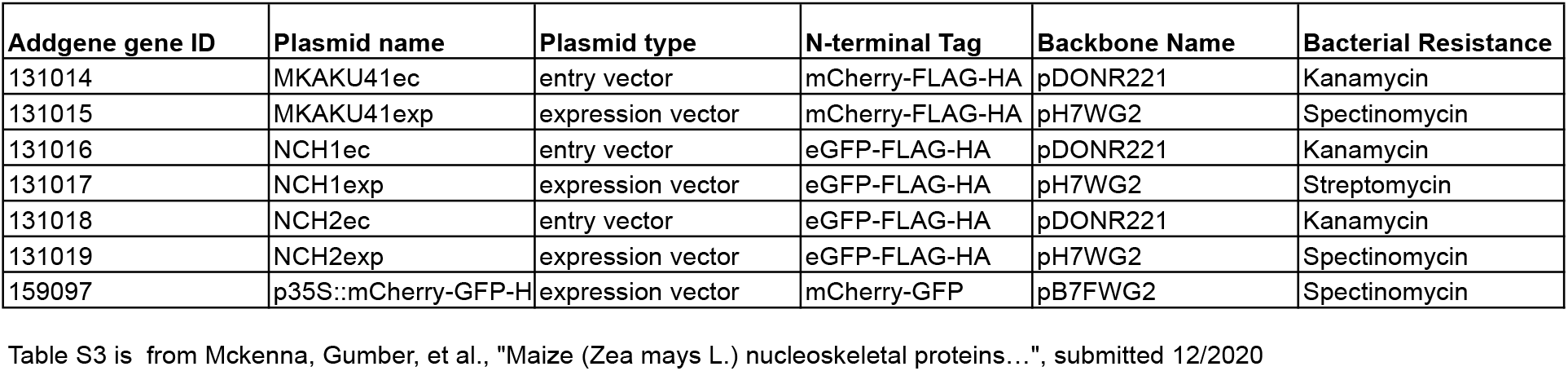
Plasmid information and Addgene IDs.

## Notes

### Competing Interest Statement

The authors have declared no competing interest.

### Summary of Updates

Revision corrects Figure 3.

